# Antibodies against Elapidae and Viperidae snake venoms: in vivo neutralization and mechanistic insights

**DOI:** 10.1101/2023.06.16.545250

**Authors:** Amit Kumar, Zaid Kamal Madni, Shivam Chaturvedi, Dinakar M. Salunke

## Abstract

Snake envenomation results in a range of clinical sequelae, and effective therapy is yet to be discovered. Anti-snake venom antibodies are being considered as a potent strategy. We developed venom-neutralizing humanized antibody scFvs and elucidated biochemical and structural mechanisms associated with the inhibition of toxicity. Tomlinson I and J human antibody scFv libraries were screened against *Naja naja* and *Echis carinatus* venoms, and seven unique antibody scFvs were obtained. Further, specific toxins of snake venom interacting with each of these scFvs were identified, and phospholipase A2 (PLA2) was found to be prominently captured by the phage-anchored antibody scFvs. Proteomic analysis of whole venom also revealed PLA2 to be the most abundant toxin in both venom samples. The scFvs binding to PLA2 were used to perform *in vivo* survival assay using the mouse model and *in vitro* toxin inhibition assays. scFv N194, which binds to acidic PLA2, showed considerable survival in *Naja naja* venom-challenged mice and conferred up to 50% protection. A combination of two scFvs, E113 and E10, both interacting with basic PLA2, exhibited synergistically enhanced survival of 33% in *Echis carinatus* venom-challenged mice, compared to 16% survival conferred by an equal amount of individual scFvs. Furthermore, these scFvs demonstrated inhibition of venom-induced myotoxicity and hemolysis which corroborate the survival data. Structural studies highlighted possible modes of PLA2 neutralization by scFv through the engagement of CDRs with C-terminal myotoxic loop and interfacial region, which are crucial for PLA2 toxicity.

## Introduction

Snakebite is a major neglected public health and medical problem, particularly in subtropical and tropical countries. According to the WHO, about 5 million snakebites befall annually, resulting in 2.7 million snake venom poisonings and 81,000 to 138,000 deaths[1, 2]. Snake venom is a complex mixture of a plethora of toxins belonging to diverse families with interspecies and intraspecies variations in the composition. Environmental conditions, geographical location, sex, type of available prey in the habitat, etc., have a substantial role in the diversification of venom composition[3, 4], which is one of the major challenges in the development of effective therapy against snakebites[5]. Primary therapeutic agents against snake venom are polyvalent serum antibodies generated in hyperimmunized animals with individual snake venom or a mixture from major species of a particular geographical area. However, several adverse effects and limitations are associated with these antivenoms[6, 7]. Among the main concerns with conventional antivenoms include various immunological responses in the host body ranging from normal allergic reactions to severe anaphylaxis[7].

Several studies have targeted the inhibition of critical toxins of snake venoms by small molecules. For example, varespladib against phospholipase A2 (PLA2)[8, 9] and dimercaprol, 2,3-dimercapto-1-propanesulfonic acid (DMPS), dimercaptosuccinic acid (DMSA), disulfiram and marimastat, batimastat against snake venom metalloproteases (SVMP)[10–12]. However, complete neutralization could not be achieved due to the lack of such inhibitors against other key toxins. Moreover, Proteins and peptides constitute most of the dry weight in snake venom and are of major pharmacological relevance[13]. Therefore, the binding of these inhibitors with protein targets is limited due to their small size of fewer than 1000 Daltons[14, 15]. On the other hand, protein drugs, e.g., scFvs and Fab, provide more specificity and selectivity toward targets due to their large size[16]. Human scFv antibodies have an added advantage of not eliciting any adverse immunological responses[17, 18] as observed in the case of conventional antivenoms in human snake bite victims. Moreover, polyvalent antivenoms consist of antibodies specific for medically irrelevant components too, which reduce the overall efficacy. The use of a phage-displayed human antibody library for screening and production of recombinant antibodies targeted exclusively for toxic venom components holds the key to obtaining high-affinity antibodies with efficient neutralization.

In the current study, we have selected venoms of two prevalent snakes of India, *Naja naja* of the Elapidae family and *Echis carinatus* of the Viperidae family, which are known to inflict distinctive toxicities[19, 20]. Tomlinson I and J libraries were screened to obtain venom-neutralizing scFvs. We further identified specific toxins of snake venom, which are interacting with these scFvs to study the impact of scFv treatment on venom-induced pathology and mortality. PLA2 was identified as the prominent interacting antigen and also the major component of snake venom which leads to many pathological reactions e.g., haemotoxicity and myotoxicity[21]. Subsequently, it was shown that the anti-PLA2 scFvs were capable of conferring protection to the venom-challenged mice. We also discovered the scFv-mediated venom inhibition and demonstrated that these scFvs were capable of inhibiting the myotoxic and hemolytic response in mice. The interaction of scFv with PLA2 was also explored at the molecular level, which highlighted key interacting residues of PLA2 that may be masked by scFvs. Structural model of E113-PLA2 complex revealed engagement of CDR regions with reported C-terminal myotoxic loop[22–24] and interfacial region[25, 26] presenting a plausible model of scFv mediated PLA2 inhibition.

## Results

### Development of potential venom-neutralizing scFvs using phage-displayed scFv libraries

The Tomlinson I and J libraries were screened against *Naja naja* and *Echis carinatus* venoms (NnV and EcV, respectively) to enrich and obtain venom-specific phage populations. After four rounds of biopanning against NnV and EcV, Tomlinson I and J polyclonal output phages were quantified at each round, and an equal number of scFv phages (10^10^ phages/well) were used to evaluate their binding with respective whole venom by polyclonal phage ELISA (Fig. S1 A-B). The fourth-round polyclonal pool was screened against each venom leading to the selection of 768 anti-NnV and 768 anti-EcV scFv monoclonal phages for further analysis. Those phages exhibiting normalized absorbance i.e. (O.D._Whole venom_/O.D._BSA_) ≥ 10, were considered as significantly specific binders. Analysis of 768 anti-NnV and 768 anti-EcV scFv monoclonal phages in monoclonal phage ELISA revealed that 231 anti-NnV and 652 anti-EcV monoclonal scFv phages displayed positive binding as per the set cut-off, to the cognate venom (Fig. 1 A-B). The 72 top binding scFv phages against each venom were selected and purified. ELISA was then carried out in triplicates using an equal number of scFv phages (10^10^ phages/well). Significant binding with the respective venoms was displayed by 53 anti-NnV and 57 anti-EcV scFv phages showed (Fig. 1 C-D). The 50 best binding phages were selected from a pool of purified monoclonal scFv phages screened against each venom, based on normalized absorbance for sequencing and further characterization.

**Fig. 1.**
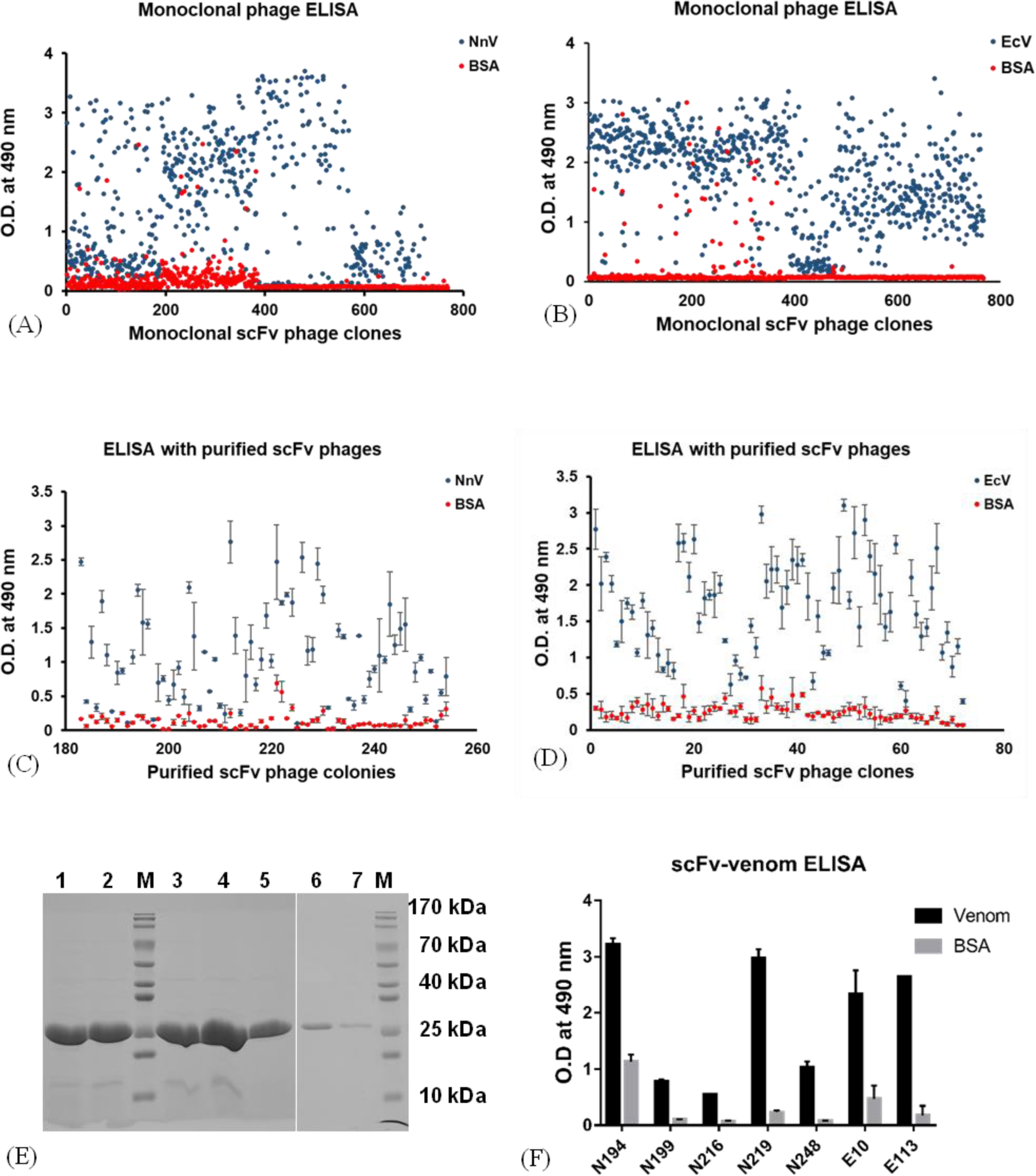
Binding of scFv clones with whole venom. The Tomlinson I and J libraries were screened against Naja naja and Echis carinatus venoms (NnV and EcV, respectively) to enrich and obtain venom-specific phage populations. (A) Binding of monoclonal scFv phages from the fourth round of biopanning with NnV. (B) Binding of monoclonal scFv phages from the fourth round of biopanning with EcV. (C) Binding of purified monoclonal scFv phages with NnV. (D) Binding of purified monoclonal scFv phages with EcV. (E) SDS PAGE gel representing purified soluble scFvs; Lane 1: N194, Lane 2: N199, Lane 3: N216, Lane 4: N219, Lane 5: N248, Lane 6: E10, Lane 7: E113, Lane M: Marker (F) Binding of soluble scFvs with the whole venom of E. carinatus and N. naja BSA was used as the negative control. P<0.05 when absorbance of venom is compared with BSA for all scFvs. In graphs C, D, and F, each data point represents mean of triplicates and error bars indicate standard deviations from mean of each data set.

Five anti-NnV (N194, N199, N216, N219, and N248) and two anti-EcV (E10 and E113) unique clones were obtained after the sequence analysis of top binding 50 anti-NnV and 50 anti-EcV scFv phages. All unique scFv clones except N199 exhibited amber stop codons in the CDR H2 region, which were replaced by glutamine. All scFvs were expressed and purified from periplasmic space. The peaks corresponding to monomeric forms of scFvs in SEC profiles were used for further characterization (Fig. S2 A-G & 1E). The purified scFvs were first tested for their binding with cognate whole venom using the direct ELISA. The N194 showed the highest binding to NnV among all scFvs, followed by N219, N248, N199 and N216. (Fig. 1F). The E10 and E113 both exhibited comparable binding to EcV. It may be noted that the binding strengths of scFvs with whole venom do not necessarily indicate binding affinity with specific interacting antigen(s) in the whole venom.

### Identification of the scFv interacting venom components

After binding analysis of scFvs with whole venom, it was imperative to identify the corresponding antigens of these scFvs for further characterization of their therapeutic role. The specific antigens in the venom corresponding to each of the scFvs were identified through pull-down assays. We used His-tag based pull-down assay to isolate the scFv-toxin complex using cobalt resin. Elutes from lanes of anti-NnV scFvs, N194, N216, N219, and N248 contained bands between 10 kDa and 15 kDa markers (Fig. S3A, S3 C-E). Bands were visible in the silver-stained gel of N199, near 50 kDa and 75 kDa (Fig. S3B). Similarly, two prominent bands near 25 kDa and 10-15 kDa (corresponding to scFv and EcV toxin, respectively) were observed in the case of anti-EcV scFvs elute of the pull-down assay (Fig. S3 F-G). Five washes were introduced before elution to avoid any contamination from non-specifically bound proteins (Fig. S3 A-G). No bands were observed in the elutes of NnV and EcV negative control (no scFvs were used), eliminating any possibility of non-specific binding of venom proteins with cobalt resin (Fig. S3 H-I).

Samples eluted from the pull-down assay were subjected to mass-spectrometry for the identification of venom components interacting with scFvs. Three filters were applied to mass spectrometry data for identification of top hits i.e., molecular weight (based on the position of toxin band(s) observed on silver-stained gels), relative abundance (in elute sample), and unique peptides. Mass spectrometry analyses of eluted samples showed N194, N216, and N248 interacted with acidic PLA2. On the other hand, N219 interacted with venom nerve growth factor (VNGF). Two bands observed corresponding to N199 were L-amino acid oxidase (LAAO) (molecular weight = 57.9 kDa) and snake venom metallo proteinase (SVMP) (molecular weight = 67.6 – 69.1 kDa) (Table 1). Pull down assays followed by mass spectrometry analysis revealed that most of the anti-NnV scFvs were binding to acidic PLA2. In fact, the critical role played by PLA2 in NnV toxicity is known[21]. Top hits in the case of E10 and E113 correspond to basic PLA2, which is also considered a major contributor towards EcV toxicity[21] (Table 1). It is noteworthy that 5 out of 7 selected scFvs interacted with PLA2.

**Table 1.**
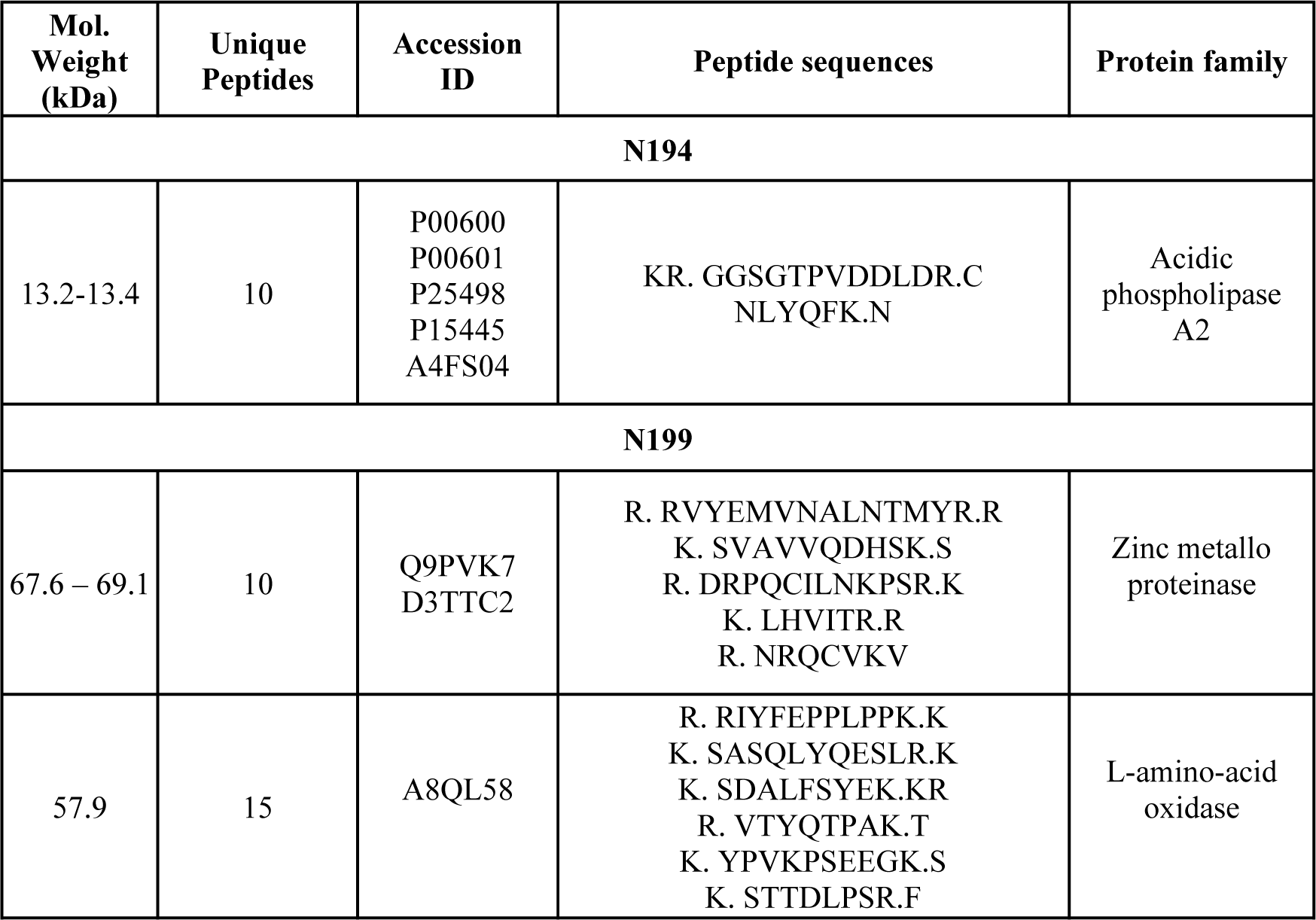
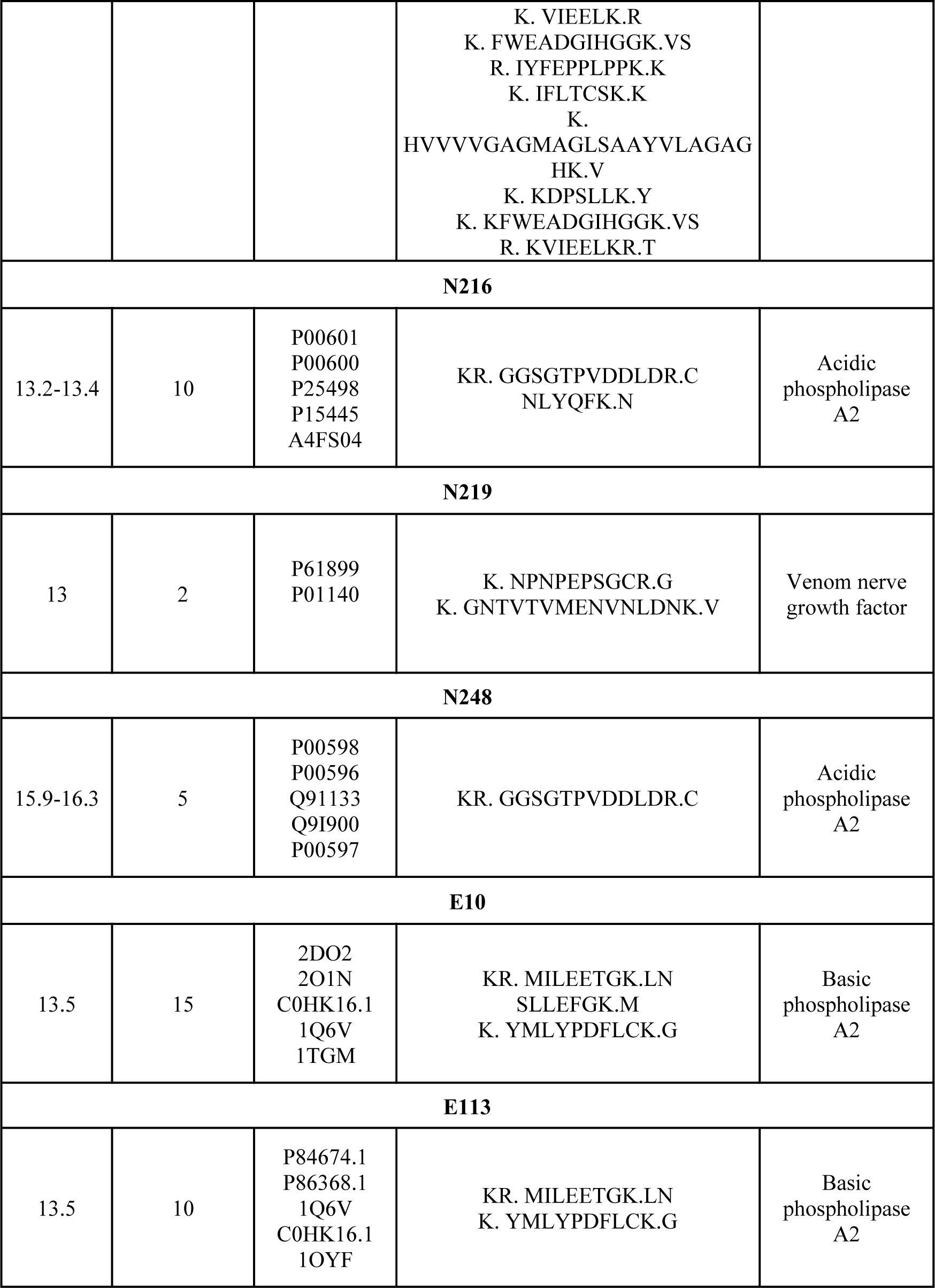
Identification of scFv interacting toxins from the whole venom. Mass spectrometry analysis of elute 1 (E1) obtained from pull-down experiment

The proteomic analysis of whole venom composition revealed PLA2 as the dominant toxin in NnV and EcV samples, with relative abundance of 24% and 28%, respectively (Fig. S4 A-D). Individual estimation of each PLA2 revealed that acidic PLA2 and basic PLA2 were dominant types of PLA2s in NnV and EcV, respectively, which is consistent with the findings of pull-down assays and explain the reason of PLA2 being the interacting antigen in the case of majority of scFvs. (Fig. S4 E-F). Cysteine-rich venom protein (CRVP), L-amino-acid oxidase (LAAO), and phospholipase B (PLB), and Three-finger toxin (3FTx) were other major components of NnV, with relative abundance of 20%, 18%, 14% and 10%, respectively (Fig. S4 A-B). Similarly, proteomic characterization of the whole venom of *Echis carinatus* revealed CRVP, snake venom serine proteases (SVSP), and LAAO as other significant toxins and accounted for 20%, 19% and 17% of the whole venom, respectively (Fig. S4 C-D). Most of the scFvs interacted with dominantly abundant toxins in their cognate venom, except for N199 which exhibited interaction with zinc metalloproteases along with abundantly present LAAO and N216 which interacted with venom nerve growth factor.

N194, N248, E10 and E113 were selected for further venom neutralization experiments, as these scFvs were interacting with crucial toxins and displayed significant binding with whole venom in ELISA. Despite of their interaction with highly abundant and very crucial toxins, N199 and N216 were not characterized further due to their low binding strength with whole venom observed in ELISA (Fig. 1F and Table 1). On the other hand, N219 was not studied further as it was interacting with not particularly abundant (Fig. S4A) and non-toxic venom component.

### Neutralization of snake venom by anti-PLA2 scFvs

Optimum doses of both venoms, anti-venom sera and selected scFvs were evaluated to perform *in vivo* survival assays. The minimum lethal doses (MLDs) of NnV and EcV were estimated to be 0.35 mg/kg and 0.55 mg/kg of mice body weight, respectively (Fig. S5). The NnV and EcV immunized mice sera were used as a positive control during *in vivo* survival assay. The binding of serum samples collected after every immunization event was evaluated (Fig. S6 A-B). Serum sample S4, collected after the third booster immunization, was selected as a positive control in the survival assay. The optimum dose of scFvs was evaluated for *in vivo* survival assay by using 2 mg and 4 mg of scFv doses E10 and E113 in EcV challenged mice, based on previous studies[41, 42] and a 4 mg scFv dose was found to be deleterious to the survival of mice (Fig. S7). Therefore, 0.5, 1, and 2 mg of scFv doses were used for further survival assays against NnV and EcV.

The antibody scFvs N194 and N248 were selected for *in vivo* survival experiments against NnV envenomation. All the mice injected with 2×(MLD) of NnV (Venom-only control group) succumbed to death within 3.5 hours. An increase in survival time was observed in all the groups treated with different doses of scFvs, up to different time intervals. Maximum protection with 50% survival till 48 hours was observed at the dose of 0.5 mg N194. In this group, the first and second mice died within 2 and 4 hours, respectively. The third mouse survived till 14 hours. The remaining 3 mice survived for 24 hours (Fig. 2A). Mice injected with 1 and 2 mg N194 too, exhibited a substantial increase in survival time. The last mouse from both groups survived for nearly 11 hours i.e., 7.5 hours more than the venom-only control group (Fig. 2A). Although N248 also conferred protection against NnV, survival rates were not comparable to those in the case of N194 treated groups. Survival was prolonged in every treated group, except in the case of group 8. About 16% survival was observed in all groups treated by N248 till 24 hours (Fig. 2B).

**Fig. 2.**
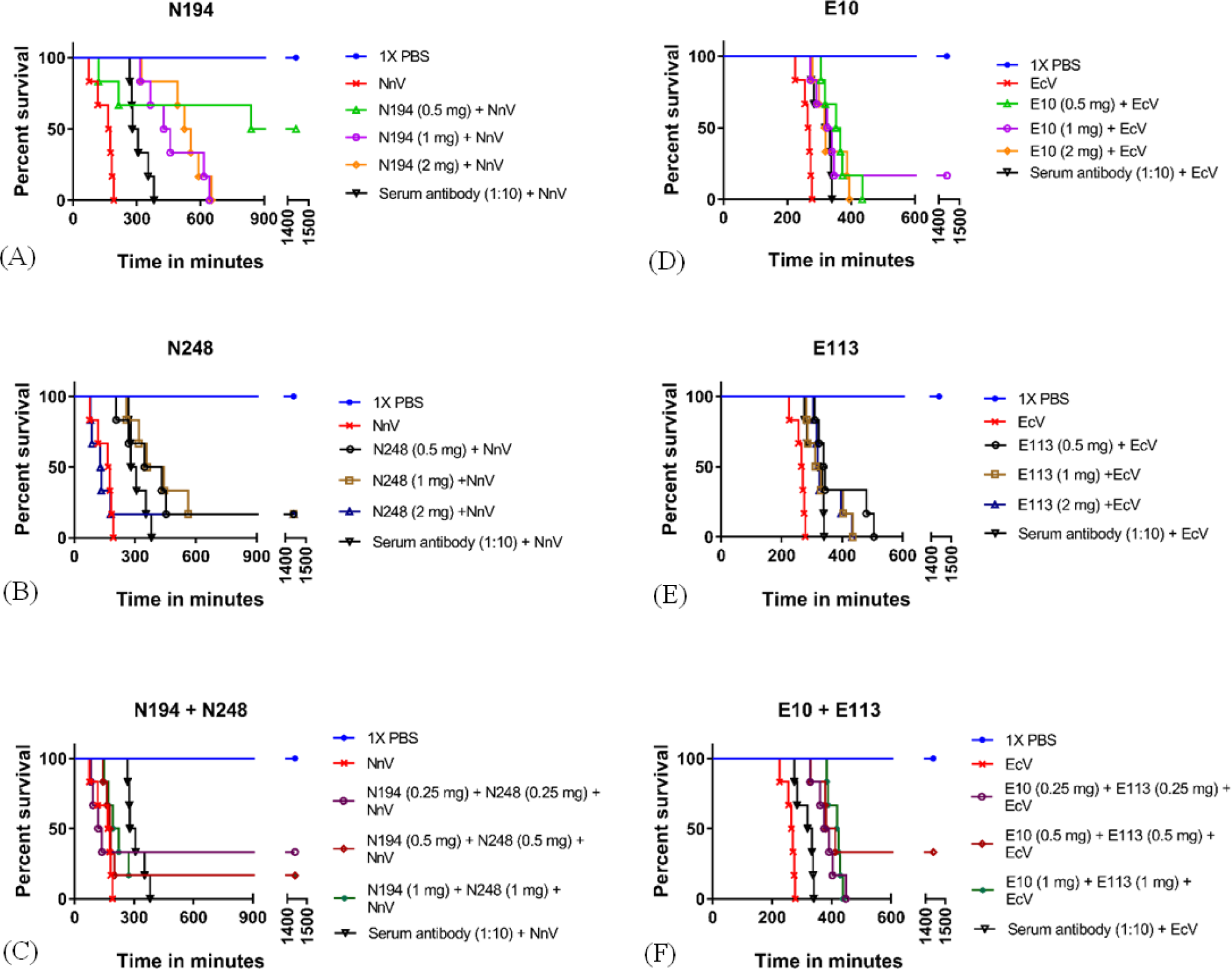
Anti-PLA2 scFvs mediated in vivo protection from the whole venom of Naja naja and Echis carinatus. Different groups of mice were intraperitoneally administered with 2xMLD of NnV (A, B, and C) and EcV (C, D, and E). Survival effects of respective scFvs were evaluated in mice groups which are given an intraperitoneal injection of the pre-incubated mixtures of 2xMLD of whole venom and different doses of scFvs. Kaplan-Meier survival analysis in (A) N194 treated group. (B) N248 treated group. (C) The group with co-administration of N194 and N248. (Vehicle control, venom-only control and serum antibody control groups were the same for all three graphs A, B, and C. Survival experiments pertaining to graphs A, B and C were performed simultaneously.) (D) E10 treated group. (E) E113 treated group. (F) The group with co-administration of E10 and E113. (Vehicle control, venom-only control, and serum antibody control groups were the same for all three graphs D, E, and F. Survival experiments pertaining to graphs D, E, and F were performed simultaneously.) All surviving mice were maintained and observed for 30 days. Each individual curve indicates the survival status of a group of 6 mice. p<0.05 when venom control groups were compared with antibody-treated groups.

To assess possible cooperativity between anti-NnV scFvs, three groups were treated with 0.5, 1, and 2 mg doses comprising an equal amount of N194 and N248. As evident from survival experiments, no indication of any cooperativity effect was observed. Combinatorial treatment with 0.25 mg of each scFv i.e., N194 and N248, resulted in 50% survival. Mice administered with higher doses, comprised of an equal amount of both scFvs, displayed 16% survival. An increase in survival time was also marginal in comparison to groups treated with N194 and N248, individually (Fig. 2A - C).

We also performed *in vivo* survival assay using E10 and E113 in EcV challenged mice. All mice in the venom control group succumbed in less than 5 hours after injection. Among E10 treated groups, the best outcome with 16% survival was conferred by 1 mg E10. Although 5 out of 6 mice died in this group, a substantial increase in survival time was observed, compared to the venom-only control group. Other E10 treated groups did not display any survival, yet significant prolongation of life was observed (Fig. 2D). The optimum dose against EcV was found to be 0.5 mg for E113, which exhibited maximum prolongation of survival time by more than 8 hours. Other doses of E113 resulted in the mortalities of all six mice within 7 hours of injection (Fig. 2E). Co-administration of 0.5 mg of each scFvs conferred maximum survival (about 33%) in mice challenged with twice the MLD of EcV among all test groups (Fig. 2F). Interestingly, these scFvs could not provide a comparable extent of protection when used individually with the same dose.

### Biochemical mechanism of protection against snake venoms

The multiple functional manifestations of PLA2 e.g., haemolysis, neurotoxicity and myotoxicity[21] provide biochemical markers for analysing the effects of antibody scFv binding with PLA2 revealing further insights into their venom neutralization potential. The effects of scFvs N194, N248, E10, and E113 on the PLA2-induced myotoxicity and hemolysis were investigated.

Myotoxicity inhibition assay was carried out by estimating Creatine Kinase activity (a known indicator of muscle damage) in the serum sample collected from mice injected with whole venom in gastrocnemius muscle[43]. N194 and N248 demonstrated almost equal inhibition in myotoxicity inflicted by NnV in a dose-dependent manner between 0.2 mg to 0.6 mg dose and displayed saturation afterward. Creatine Kinase activity was reduced from 568 units/L (venom control group) to 255.8 and 246.3 units/L upon administration of the N194 dose of 0.6 mg and 0.8 mg, respectively (Fig. 3A). Similarly, treatment with 0.6 mg and 0.8 mg of N248 resulted in the decline of Creatine Kinase activity from 568 units/L to 249.7 and 217.7 units/L, respectively (Fig. 3B). Myotoxicity was reduced by more than 2-fold at higher concentrations of scFvs in comparison to the venom control group (Fig. 3 A-B).

**Fig. 3.**
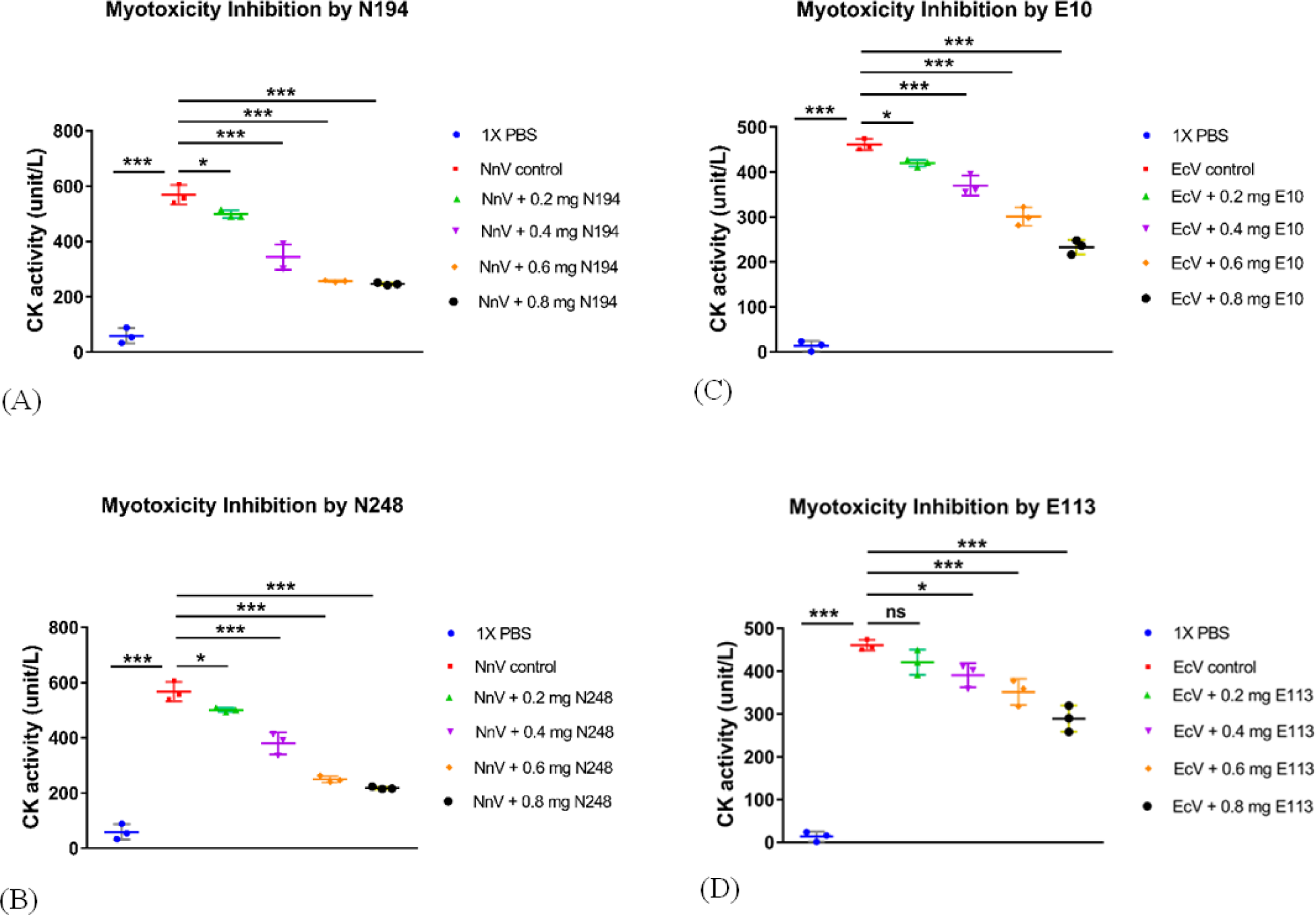
Evaluation of venom induced myotoxicity inhibition by anti-PLA2 scFvs. Myotoxicity inhibition was evaluated by estimating Creatine Kinase activity in the serum sample of mice injected (intramuscular injection in gastrocnemius muscle) with pre-incubated mixtures of the whole venom and different doses of scFvs. Myotoxicity inhibition in (A) N194 treated group. (B) N248 treated group. (Vehicle control and venom-only control groups were the same for graphs A and B. Myotoxicity inhibition assays pertaining to graphs A and B were performed simultaneously.) (C) E10 treated group. (D) E113 treated group. (Vehicle control and venom-only control groups were the same for graphs C and D. Myotoxicity inhibition assays pertaining to graphs C and D were performed simultaneously.) Each data point represents the mean of triplicates, and error bars indicate standard deviations from the mean of each data set. For all the graphs n=3 and p<0.05, except the group treated with 0.2 mg of E113(ns-non-significant, *p=0.03, **p= 0.002, ***p<0.001).

During EcV induced myotoxicity assay, Creatine Kinase levels of 13.47 units/L and 460.67 units/L were calculated in the vehicle control group and venom control group, respectively. E10 and E113 demonstrated dose-dependent inhibition in myotoxicity. A two-fold reduction in myotoxicity was observed at a 0.8 mg dose of E10, with a Creatine Kinase level of 233.22 units/L. A minor decrease in Creatine Kinase level from 460.67 units/L to 419.56 units/L was observed in a lower E10 dose of 0.2 mg, followed by 370.13 units/L, 300.96 units/L and 233.21 units/L with a corresponding scFv dose of 0.4 mg, 0.6 mg, and 0.8 mg, respectively (Fig. 3C). Likewise, incubation with E113 also resulted in a reduction in myotoxicity but up to a lesser extent compared to E10. The Creatine Kinase levels of 420.7 units/L, 390.25 units/L, 351.44 units/L, and 288.91 units/L were calculated for cognate E113 doses of 0.2 mg, 0.4 mg, 0.6 mg, and 0.8 mg, respectively (Fig. 3D). Although both scFvs inhibited EcV-induced myotoxicity in mice, E10 was found to be a stronger inhibitor than E113. The myotoxicity inhibition potential of E10 and E113 explained the mechanism of survival conferred by these scFvs (Fig. 3 C-D).

The effect of anti-NnV and anti-EcV scFvs on venom-induced haemolysis was also analyzed by estimating released haemoglobin at 415 nm. E113 reduced EcV-induced haemolysis in a dose-dependent manner, and up to 50% of haemolysis inhibition was achieved at 2 mg/ml scFv concentration (Fig. 4D). However, N194, N248, and E10 failed to inhibit haemolysis induced by their respective venoms (Fig. 4 A-C). N194 and N248 exhibited a comparable extent of reduction in myotoxicity, yet protection against NnV conferred by N194 was substantially higher than that conferred by N248 (Fig. 2 A-B & 3 A-B). N194 might be involved in the inhibition of other roles of acidic PLA2 e.g., pre-synaptic neurotoxicity, platelet aggregation, etc., which can account for an increased survival rate[21, 44, 45]. On the other hand, enhanced survival from EcV envenomation during combinatorial treatment by E10 and E113 can be accounted for by the dual function of E113 mediated inhibition of hemotoxicity and myotoxicity, along with stronger inhibition of myotoxicity by E10 (Fig. 2F, 3 C-D & 4F).

**Fig. 4.**
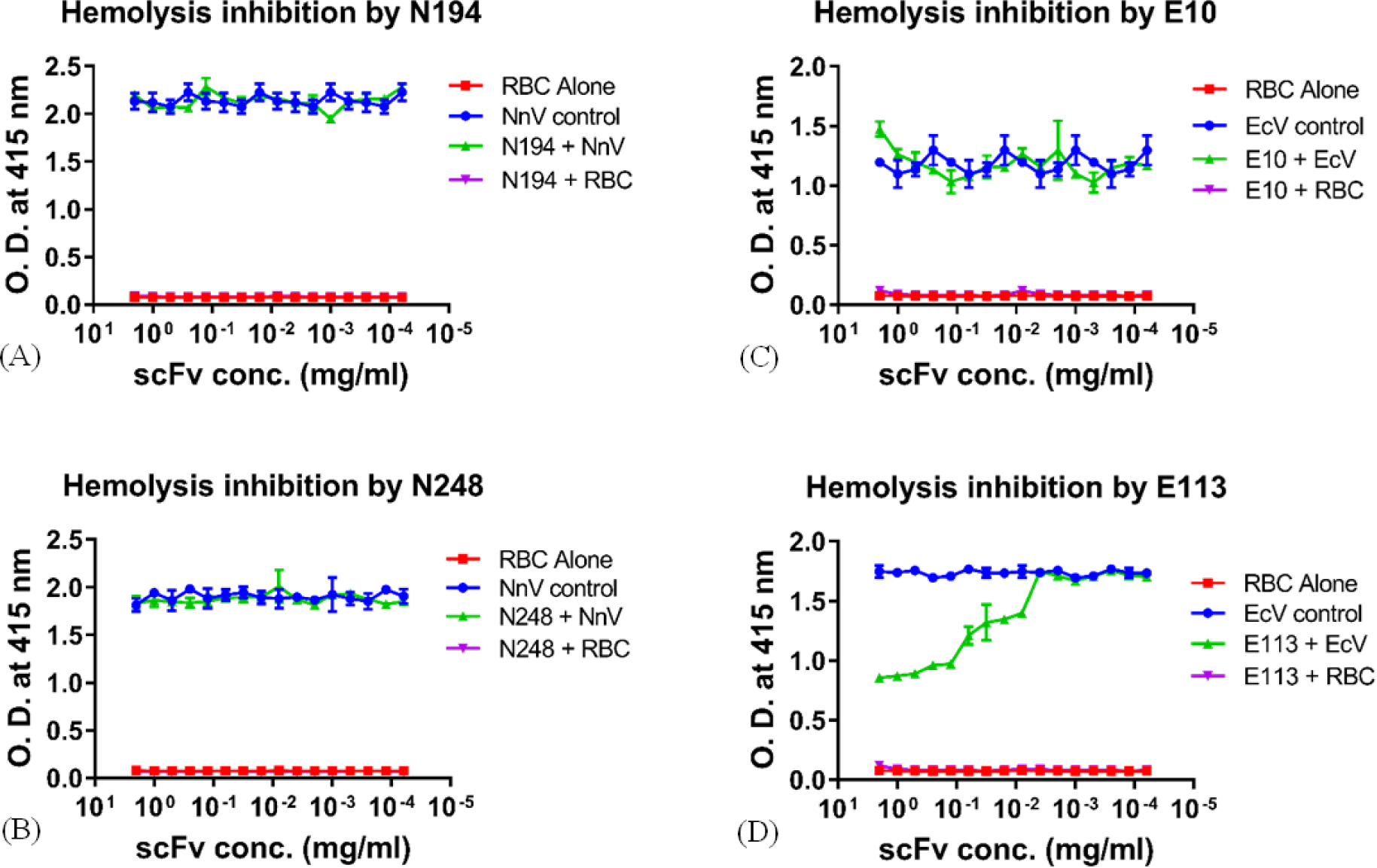
Evaluation of venom-induced haemolysis inhibition by anti-PLA2 scFvs. (A) N194 treated group. (B) N248 treated group. (C) E10 treated group. (D) E113 treated group. Each data point represents the mean of triplicates, and error bars indicate standard deviations from the mean of each data set. For graphs D, p<0.05.

### Structural understanding of scFv-mediated PLA2 inhibition

Towards understanding the structural basis of PLA2 inhibition crystallization attempts were made involving the scFvs which resulted in good quality crystals of E113. The crystal structure of E113 was determined at 2.75 Å resolution using molecular replacement method (Fig. 5A). To understand molecular interactions between E113 and PLA2 and their role in the inhibition of the myotoxic effect of basic PLA2, molecular docking was carried out using crystal structure of E113 and the reported structure of PLA2 (PDB ID: 2QHE). Basic PLA2 consists of three α-helices viz., N-terminal α-helix, two antiparallel α-helices, two antiparallel β-sheets, one putative Ca^2+^ binding loop and C-terminal myotoxic loop[22]. The suramin is a known strong inhibitor of myotoxicity induced by PLA2[22, 25]. Structural details of PLA2/suramin complex (PDB ID: 3BJW) highlighted some key interactions of suramin with C-terminal loop (K115-K129) of PLA2, suggesting a crucial role in myotoxicity[22]. Role of C-terminal loop (K115 - K129) has also been established through other studies[23, 24, 26]. In addition, the significance of i-face (formed by R34, D53, and K69) in binding with inhibitors has also been studied[25].

**Fig. 5.**
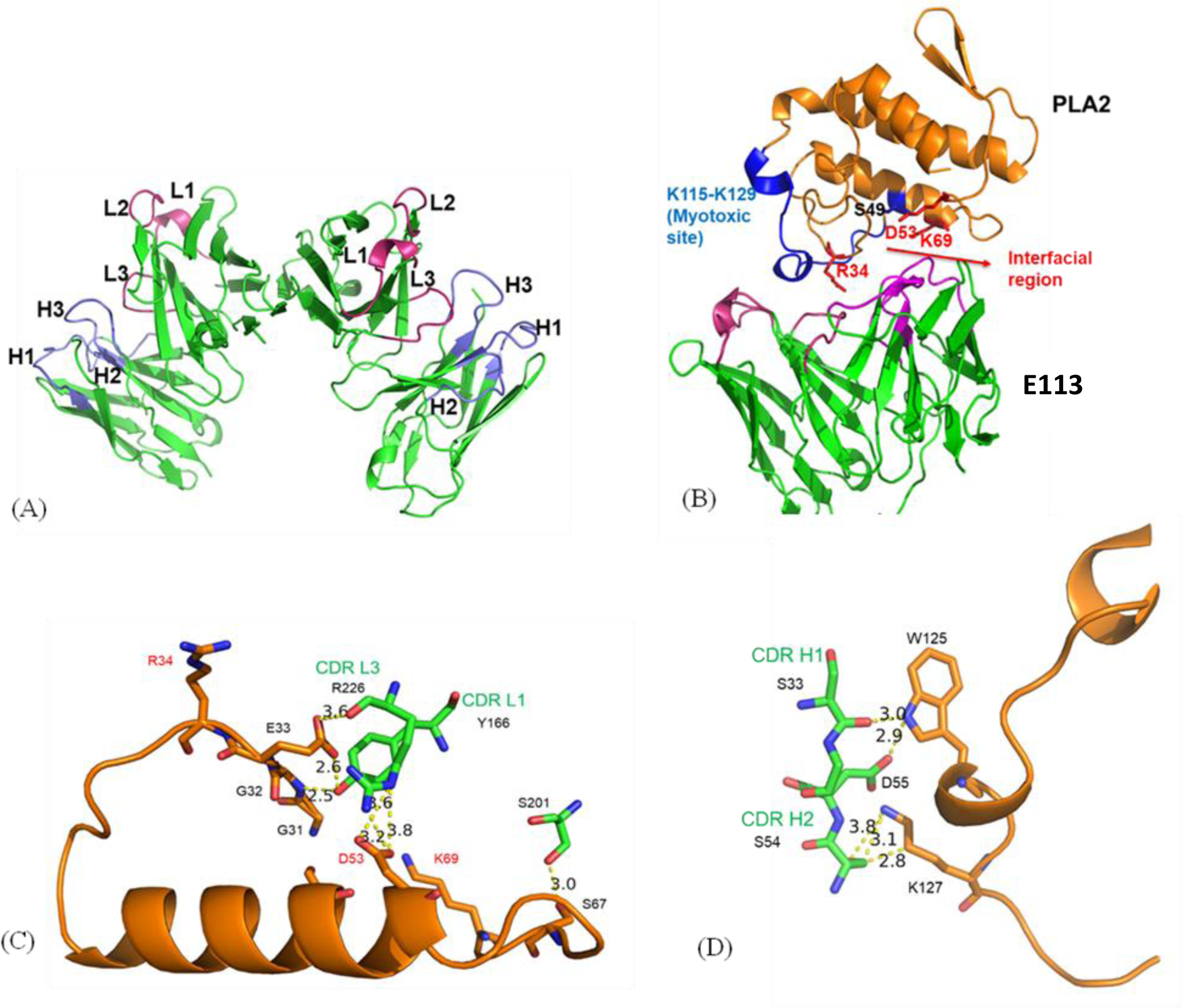
Structural basis of PLA2 inhibition by E113. (A) Crystal structure of E113 with two molecules in one asymmetric unit (PDB ID: 8IA6). (B) Interaction of E113 with PLA2. (Myotoxic loop is shown in blue colour. Residues comprising i-face, are shown in red colour.) (C) Interaction of light chain of E113 with i-face of PLA2. (D) The interaction of heavy chain of E113 with the myotoxic loop of PLA2.E113 is shown in green. PLA2 is shown in orange.

The E113 was docked on the PLA2 (PDB ID: 2QHE) using the Schrödinger suite. The best docked pose was selected based on the proximity of CDRs of scFv and myotoxic loop and i-face of PLA2, with the *in silico* relative binding affinity of 27.70 kcal/mol (Fig. 5B). To examine stability of the complex E113/PLA2, an MD simulation run was carried out for 500 ns, and no major structural deviations were observed among the snapshots of the complex extracted at different time frames (Fig. S10A). The root mean square deviation (RMSD) and the root mean square fluctuation (RMSF) of the whole complex was found to be below 4.5 Å and 2.5 Å, respectively. No significant fluctuations were observed during the simulation run, except for residues corresponding to flexible Gly-Ser linker connecting heavy chain with light chain (Fig. S10 B-C).

It was observed that CDR L1 and CDR L2 interacted with the i-face region, which is the recognition site of suramin (Fig. 5B). Y166 (CDR L1) interacted with G32 and E33 mainly by hydrogen bonds (H-bonds). H-bonds were also observed between R226 (CDR L3) and E33. R226 (CDR L3) formed H-bonds with D53. In addition, S201 of E113 formed H-bond with S67. It was evident that E113 interacted with the i-face region by establishing contacts with D53 and neighbouring residues (Fig. 5C). E113 also interacted with the myotoxic loop through CDR H1 and CDR H2 (Fig. 5B). S33 from CDR H1 interacted with W125, by several H-bonds. S54 (CDR H2) interacted with W125 and K127 of the myotoxic loop, which are important residues involved in myotoxicity (Fig. 5D). Thus, the E113/PLA2 complex model revealed that CDRs encompassed crucial regions of PLA2 i.e., i-face and c-terminal myotoxic loop. Structural studies of scFv mediated PLA2 inhibition indicated that the binding of scFv with these crucial regions blocked the interaction of PLA2 and interfered with its activity (Fig. 5 B-D).

## Discussion

Conventional serum-based antivenoms are the major available therapies against snakebite even though the manufacturing processes are complex and they are associated with many side effects[1, 5, 46]. Also, huge diversity and extensive heterogeneity in venom composition are among the hurdles in antivenom development[5, 47]. However, in recent decades human therapeutic antibodies and their relevant domains are showing potential as effective biologics for the treatment of various diseases[48]. This approach is also being explored towards generating therapeutic antibodies against snake bite[17, 41, 42, 49–54]. In current study, we obtained four venom neutralizing scFv antibodies which displayed *in vivo* and *in vitro* venom inhibition. Biochemical and structural basis of the antibody scFv effect on venom toxicity was also deciphered.

Antibody phage display approach is a promising tool to develop therapeutic monoclonal antibodies as the phage-displayed antibody technology mimics the immune system *in vitro*[55, 56]. We characterized a vast number of monoclonal scFv phages after screening the antibody scFv phage display library. The sequence analysis of the top binding monoclonal phages for each venom revealed the presence of redundant sequences in the selected phage populations, which can be attributed to more number of biopanning rounds and selection of monoclonal phage populations from final screening round[51, 52, 57, 58]. Thus, the lower specificity of the polyclonal phage population can be correlated to higher sequence diversity in the monoclonal phage population during the screening of the phage-displayed antibody library using the biopanning process.

Most of the anti-NnV and anti-EcV scFvs were found to be interacting with acidic and basis PLA2, respectively. This corroborates with proteomics analysis of whole venom, identifying PLA2 to be the most abundant toxin in both the venoms. The distribution of PLA2 varies significantly among the same snake species of *E. carinatus* from distinct geographical regions[4, 10, 12]. Other crucial toxins in the whole venom of *Echis* species vary substantially in terms of the extent of abundance[4, 10, 12]. Similar diversity in toxin composition has been reported in *N. naja* venom obtained from other geographical areas[4]. Two of the anti-NnV antibody scFvs, N194 and N248, against *N. naja* and another two anti-EcV scFvs, E10 and E113, against *E. carinatus* were binding well to PLA2 and were selected for venom neutralization studies. The scFv N199 was found to be interacting with two toxins, the relatively abundant LAAO, and comparatively less abundant SVMP. N216 binds to VNGF, which is less commonly observed in venoms. Monoclonal scFvs didn’t interact with other dominant toxins of both venoms e.g., CRVP, PLB, 3FTx, SVSP, and LAAO and others.

The survival assays for assessment of the neutralizing potential of selected scFvs against respective venoms showed that N194 conferred significantly higher survival than N248, which was consistent with the difference in their binding with NnV. On the other hand, co-administration of E10 and E113 resulted in synergistically enhanced survival of EcV envenomed mice, compared to the group injected with an equivalent dose of individual scFvs. The survival percentage observed in the current study is comparable to other independent survival studies[10, 53]. In a similar study, treatment with a cocktail comprising two and three mAbs resulted in substantial enhancement in survival than single mAb, in mice challenged with *Dendroaspis polylepis* venom fraction[53]. In another study, only combinatorial treatment with varespladib, nafamostst, and marimastat, resulted in effective neutralization of *E. ocellatus* and other viper venoms[10]. It was noteworthy that in the current study, maximum protection was observed at a specific optimum concentration of each scFv.

Myotoxicity is a characteristic feature of most Viperidae and some Elapidae snake envenomations. PLA2 is a vital toxin for the induction of tissue damage by damaging the structural integrity of muscle fiber plasma membrane[59–61]. While many venom components contribute to hemotoxicity, PLA2 is the major toxin causing hemolysis[62–64]. The survival against snake venom conferred by different anti-PLA2 scFvs and their combinations were also manifested in terms of the effect on PLA2-associated pathologies. The effect of each scFv on venom-induced inhibition of hemolysis was observed and only scFv E113 reduced it significantly. Although the other three scFvs did not show any impact on hemolysis, a significant role in venom neutralization was observed during *in vivo* survival assay. This indicated the possible role of these scFvs in a different mode of PLA2 inhibition. All four scFvs displayed myotoxicity inhibition to varied extents. In the case of anti-NnV scFvs, both scFvs reduced NnV-induced myotoxicity by almost 50%. Almost equal reduction in myotoxicity inhibition was conferred by both scFvs. However, N194 demonstrated significantly higher survival than N248 in the mice model, which indicated the possibility of an N194 role in the abrogation of other PLA2-associated pathologies[21]. Likewise, both anti-EcV scFvs, E10 and E113, exhibited inhibition in EcV-induced myotoxicity, but the extent of inhibition was lower, and saturation in inhibition was not observed even at higher concentrations of scFvs. E10 was found to be a slightly more potent myotoxic inhibitor, compared to E113. Enhanced survival observed in groups co-administered with these scFvs, could be attributed to simultaneous stronger myotoxic inhibition by E10 and inhibition in haemotoxicity by E113. The benefits of combinatorial therapy were established by observations made during survival and toxin inhibition assay, which could be attributed to inhibition of EcV-induced RBC lysis by E113 in addition to myotoxicity inhibition by both anti-EcV scFvs. The use of an adequate proportion of scFvs targeting different crucial toxins or even the same toxins but inhibiting different pathologies could yield enhanced therapeutic outcomes.

The structural basis of venom neutralization using a model based on the crystal structure of E113 and the already reported structure of *E. carinatus* PLA2 was analyzed. E113 was found to interact with key residues of the myotoxic loop and interfacial region of PLA2. These interactions are similar to those observed at the binding site of suramin, a potent PLA2 inhibitor[22, 25]. This suggested that E113 is likely to adopt a similar mechanism as that by suramin to abate myotoxic activity. Although this study presents a plausible structural basis for PLA2 inhibition, the crystal structure of the E113/PLA2 complex would give a more concrete explanation of scFv-mediated PLA2 inhibition. It is possible that the scFvs screened against other crucial venom toxins might also contribute to snake venom neutralization. The structural insights obtained here concerning the mode of PLA2 inhibition by scFv can be used to optimize and re-engineer therapeutic antibodies by selective mutations in key residues.

The next-generation antivenoms will be comprised of oligoclonal mixtures of antibodies targeting all medically relevant toxins that could provide an effective solution to the present menace of snakebite[53, 18]. Of course, various factors need to be taken into consideration while developing recombinant antivenoms. Loss of diversity in the scFv pool during the panning round, amplification of coat protein-specific scFvs, and masking of antigen active sites are some of the major challenges associated with antibody phage display technology[49, 65]. Moreover, proportions of scFvs should be in accordance with toxicity score and abundance of respective toxins for effective neutralization[18, 66]. Five out of seven unique scFvs were interacting with PLA2, the most abundant venom component in the two snake varieties explored, and four anti-PLA2 scFvs conferred *in vivo* protection against whole venom. However, the protective efficacy of these scFvs in the mouse model could also be limited due to their human origin. Structural insights into scFv-toxin interaction are crucial for redesigning therapeutic antibodies by selective mutations in participating residues or by the introduction of the additional fragment to achieve improved affinity, more stability, and less autoreactivity[67–69].

## Materials and methods

### Animals and Biological products

Whole venom of *Echis carinatus* (EcV) and *Naja naja* (NnV) were procured from Irula Snake Catchers Industrial Co-operative Society, Tamil Nadu, India. Male and female BALB/c mice were bred in institutional animal house. Clearance for use of the animal model was taken from the Institutional Animal Ethics Committee (IAEC) of ICGEB, New Delhi, and IAEC approval no. is ICGEB/IAEC/07032020/SI-17. While using animal models, internationally accepted guidelines and regulations for experimental use and care were followed.

### Antibody phage display screening

The screening procedure of Tomlinson I and J libraries (Medical Research Council, Centre for Protein Engineering (MRC-CPE), Cambridge, United Kingdom Resource Centre, Cambridge, UK) was adopted from manufacturer’s protocol. Briefly, MaxiSorp and PolySorp immunotubes (Nunc, Thermo Fisher Scientific, USA) (in every alternate round) were immobilized with 4 ml of 100 μg/mL whole venom and BSA in 1X PBS and kept at 4°C overnight. Thereafter, the tubes were washed with PBS thrice. Blocking solution of either 2% skim milk (Hi-Media Laboratories Ltd., India) or protein free blocking solution (Geno Technology Inc., USA) was used every other cycle for 2 hr at 25°C. The Tomlinson I & J libraries were afterward incubated to the whole venom coated immunotubes for 1 hr on rotating platform and 1 hr static incubation. The unbound phages were eliminated by aspiration and immunotubes were washed with 1X PBS with 0.05% Tween 20 (SigmaAldrich Corp., USA) (10 times for selection round 1 and incremented 10 times after every subsequent round) to remove the weakly bound phages. Affinity elution was done with 0.45 mL of 1 mg/ml of whole venom for 30 min to elute the bound phages. The eluted phages were then treated with 50 μl of bovine trypsin (SigmaAldrich Corp., USA) (10 mg/mL) for 10 min. 250 μl of eluted phages of each Tomlinson I & J libraries were infected with 1.75 ml of TG1 at an OD_600_ of 0.4 for amplification of phages and the remaining 250 μl were stored at 4°C. The infected culture was incubated at 37°C for 30 min. The 100-fold dilution of culture was made up to 10^-4^ dilutions and spotted on TYE plates (Hi-Media Laboratories Ltd., India) having 1% glucose (Merck & Co., USA) and ampicillin (Gold Biotechnology, Inc., USA). The remaining TG1 culture was spun at 11,600g and the pellet was plated on TYE plates.

The next day, the plates spotted with the pellet were scraped and used for amplification and purification of polyclonal scFv phages as per manufacturer’s protocol. The phage output obtained during the 1^st^ cycle of selection was used as input for the second cycle and similarly goes on for further rounds. The whole process is repeated for 4-5 times to get the high-affinity phages binding against the whole venom.

### Phage screening by ELISA

#### Polyclonal phage ELISA

The phages obtained after 4 rounds of selections were checked for their binding towards the EcV or NnV. 100 μg/mL of EcV or NnV and BSA was coated on a 96-well microplate. On the following day, the plate was washed thrice with 1X PBS and blocked with 2% skim milk in 1X PBS for 2 hr. 10 μl of polyclonal phages were added to the plate from every round of selection. Incubation was done for one hr at room temperature. The phages were removed and plates were washed thoroughly with 0.1% Tween 20 in PBS (PBST). The anti-mouse M13-HRP (GE Healthcare Cat# 27942101) was added at 1:5000 dilution for 1hr at room temperature. PBST was used as a washing solution for 96-well microplates and were subsequently developed with ortho-phenylenediamine (OPD) (SigmaAldrich Corp., USA) and H_2_O_2_ (SigmaAldrich Corp., USA) as a substrate. The absorbance was read at 490 nm.

#### Monoclonal phage ELISA

The individual isolated colonies from the 4^th^ round of biopanning were inoculated in 200 μl of 2×TY media (Hi-Media Laboratories Ltd., India) having ampicillin and 1% glucose in 96 well round bottom plate. The culture was grown overnight at 37°C. On the subsequent day, 20 μl of inoculum was transferred into 200 μl of 2×TY with ampicillin and 1% glucose in a fresh flat-bottom 96-well plate. The culture was grown for 2 hr at 37°C and stocks were made from the original plate by adding glycerol (Merck & Co., USA) to the final concentration of 15% and stored at -70°C. After incubation, 25 μl of 2×TY containing 10^9^ helper phages were added to 96 well plate. The plates were further incubated at 37°C for 1 hr with shaking at 200 rpm. The culture was then spun at 1800g for 10min, and the pellet was resuspended in 200 μl of 2×TY with ampicillin, kanamycin (Gold Biotechnology, Inc., USA) and grown overnight at 30°C. The culture was spun again, and the supernatant was used in ELISA using the protocol as discussed in polyclonal phage ELISA.

### Cloning, expression, and purification of scFvs

The pIT2 phagemids containing scFv genes were isolated from the 5ml of bacterial culture using the Qiagen Miniprep plasmid isolation kit and digested using the restriction enzyme *Nco* I (Thermo Fisher Scientific, USA) and *Not* I (Thermo Fisher Scientific, USA). Digested scFv inserts were subcloned in pET22b(+) vector (EMD Biosciences) and transformed into BL21(DE3) *E-coli* strain (Thermo Fisher Scientific, USA). Amber stop codon in CDR regions of scFvs were mutated to glutamine by site-directed mutagenesis kit (Thermo Fisher Scientific, USA). The primers for the SDM were made using online software PrimerX[27]. The pET22b(+) plasmids containing mutated scFvs were transformed in BL21 cells and induced with 1 mM IPTG (SigmaAldrich Corp., USA) at O.D._600_ of 1.5. The culture flasks were incubated at 18°C overnight. Harvested cells were used for periplasmic extraction of expressed scFvs by employing the method used by Christ *et al.* with slight modifications[28] and purified using a combination of immobilized metal affinity chromatography and gel filtration chromatography. The supernatant obtained after periplasmic extraction was loaded onto the pre-equilibrated column at the rate of 0.25 ml/min in the peristaltic pump at 4°C temperature. The elution was done in a linear gradient of 20-100% for 90 min with 50 mM Tris-HCl pH-8.0, and 150 mM NaCl containing 1M imidazole using FPLC. The protein sample obtained after Ni-NTA was concentrated up to 5 ml using a centrifugal filter unit (Merck & Co., USA) with a cutoff size of 10 kDa. The concentrated protein sample was loaded on a pre-equilibrated Superdex-75 16/60 column (GE Healthcare, USA). A buffer consisting of 50 mM Tris-HCl (pH 8.0), 150 mM NaCl and 10% glycerol was used for equilibrating the column as well as the running buffer. The protein purity was checked using 12% SDS-PAGE. Final purified scFvs i.e., scFv-10-α-EcV, scFv-113-α-EcV, scFv-194-α-NnV, scFv-199-α-NnV, scFv-216-α-NnV, scFv-219-α-NnV, and scFv-248-α-NnV were designated as E10, E113, N194, N199, N216, N219, and N248, respectively.

### Proteomic analysis of whole venom

The whole venom sample (25 μg) was digested using Trypsin (1:50 dilution) for overnight at 37 °C. C18 silica cartridge was used for cleaning of digest by salt removal and further dried by speed vacuum drier, followed by its reduction using 5 mM TCEP and alkylation with 50 mM iodoacetamide. Dried peptide sample was dissolved in buffer A (2% acetonitrile, 0.1% formic acid) and 1 μg of peptide sample was loaded on Acclaim PepMap 75 μm x 2 cm and C18, 3 μm particle size guard column and eluted using a gradient of buffer B (80% acetonitrile and 0.1% formic acid). Eluted peptides were separated on C18 column 15 cm, 3.0 μm Easy-spray column (Thermo Fisher Scientific, USA) at a flow rate of 300 nl/min), followed by injection for MS analysis. LC gradients were run for 60 minutes. MS1 spectra were acquired in the Orbitrap (R= 120K; AGQ target = 300%; RF Lens = 80%; mass range = 375−1500; centroid data), followed by collection of MS2 spectra for top 10 peptides. The raw data was further analyzed using Proteome Discoverer (v2.4) against the Uniprot protein databases. 10 ppm and 0.02 Da cut-off was set for the precursor and fragment mass tolerances, for Sequest search, respectively. Carbamidomethyl on cysteine was selected as fixed modification whereas N-terminal acetylation and methionine oxidation were selected as variable modifications. 0.01 FDR were set for peptide spectrum match and protein false discovery rate.

### Pull down assay

The scFv expressed and puried has His-tag at the N-terminal. 200 µg of scFvs were incubated with whole venom for 12 hours at 4°C. The scFv + venom mixtures were incubated with for 6 hours. Pre-incubated mixture of scFv and respective whole venom was mixed with high-performance His-tag binding cobalt resin beads provided with Pierce™ His Protein Interaction Pull-Down Kit (Thermo Fisher Scientific, USA) and incubated at 20°C for 12 hours to allow attachment of the complex on resin via His-tag. Flow through was collected in collection tubes by spinning the mixture for 1 minute for further analysis. Cobalt resin was washed five times with 1X TBS and 20 mM imidazole to ensure the removal of any unbound, non-specifically, or weakly bound components of scFv-venom mixture. Wash samples after 5^th^ wash were collected and stored to verify the complete removal of any unbound components. The attached scFv-toxin complexes were then eluted using 500 mM of Imidazole containing buffer after washing 5 times with 20 mM Imidazole containing buffer. The eluted samples were run on 15% SDS-polyacrylamide gel under reducing conditions followed by staining by silver staining solution using Pierce™ Silver Stain Kit (Thermo Fisher Scientific, USA) (Fig. S3 A-G). The pull-down experiment was also performed separately with EcV and NnV in the absence of scFvs and taken as a negative control (Fig. S3 H-I). Pull-down eluted samples were analysed by mass spectrometry to identify venom toxins interacting with scFvs. Top 5 hits in mass spectrometry were selected as scFv interacting antigens based on molecular weight, relative abundance, and unique peptides (highest to lowest) filters.

### *In vivo* protection assay

BALB/c female mice were used to perform the protection assay. The sample size of BALB/c mice was calculated using one-way ANOVA by G*Power 3.1.9.6 software for MLD estimation and survival experiments, with effect size = 0.47, α error probability = 0.2 and power (1-β error probability = 0.8) for MLD estimation and scFv dose optimization. Group sizes for *in vivo* were computed using same method with effect size = 0.45, α error probability = 0.2 and power (1-β error probability = 0.8).

#### Minimum lethal dose (MLD) estimation

The minimum lethal dose (MLD) was estimated using female BALB/c mice and used accordingly to perform survival experiments. For EcV MLD estimation, eight groups with six mice in each group, of age between 6-8 weeks and a weight range between 20-22 grams, were administered with saline and increasing dose of whole venom between 0.5 mg/kg and 1.5 mg/kg of mice body weight for EcV (Table. S1A). In order to estimate the minimum lethal dose of NnV, ten groups with 6 mice in each group were selected. Group 1 was taken as negative control and injected with saline. *Naja naja* whole venom was dissolved in saline and injected intraperitoneally in groups 2 to 10 with 0.2 mg/kg to 1 mg/kg of body weight, as described in table S1A. These mice were observed for 48 hours for their survival and other clinical symptoms arising due to envenomation. Lowest amount of administered venom causing 100% death within 48 hours, was considered as MLD.

#### Survival assay (for scFv dose optimization)

Nine groups, consisting 6 female BALB/c mice in each group weighing 20-22 grams, were used to estimate optimum scFv dose for *in vivo* survival assay. Saline, 3×MLD of EcV, and pre-incubated mixture of 3×MLD EcV with different scFv doses, were injected intraperitoneally as mentioned in table S1B. The number of surviving mice at the end of 8 hours was recorded for each group.

#### *In vivo* survival assay

A total of 12 groups, consisting of six BALB/c female mice in each group, were used for survival assay. All mice were 6-8 weeks old and weighed around 20-22 grams. Different doses scFvs were incubated with 2×MLD of venom at room temperature for 30 minutes and injected into mice intraperitoneally as shown in table S1C. Mice challenged with saline and venom were considered as a vehicle and venom control. The number of surviving mice at the end of 48 hours was recorded for each group. All visible clinical symptoms were also observed and recorded.

Kaplan-Meier survival graph were plotted for each group using GraphPad PRISM 7 software, to estimate survival percentage.

### Myotoxicity inhibition assay

BALB/c male mice were used to perform the myotoxicity inhibition assay. The sample size of BALB/c mice was calculated using one-way ANOVA by G*Power 3.1.9.6 software as described earlier, with effect size = 0.48, α error probability = 0.2 and power (1-β error probability = 0.8).

During NnV and EcV optimum dose estimation for myotoxicity inhibition assay, 5 groups of 5 mice in each group between weight range of 20-22 grams, were injected with saline (negative control) and different amounts of venom in the Gastrocnemius muscle of the right leg, as mentioned in table S2A. All mice injected with 50 μg of NnV, succumbed to death within 2 hours, so Creatine Kinase activity couldn’t be estimated for this group. After incubation for 2 hours, 100 µl of blood was collected from each mouse and kept for 1 hour at 37°C. Blood samples were then centrifuged at 3000 rpm for 30 minutes at 4°C. Serum samples were collected from surviving mice and 3 samples from each group were used for estimation of Creatine Kinase activity Assay Kit (Sigma-Aldrich).

110 µl of water and 10 µl of calibrator buffer in 100 µl of water were used as blank and calibrator, respectively. 10 µl of serum samples were mixed with 100 µl of reconstituted buffer. All samples were incubated at 37°C for 20 minutes. (A_340_) initial and (A_340_) final were calculated as absorbance of test samples measured at 340 nm after 20 minutes and 25 minutes, respectively. Similarly, the (A_340_) blank and (A_340_) calibrator were also calculated after 25 minutes. Creatine Kinase activity was finally measured by using the following formulae.

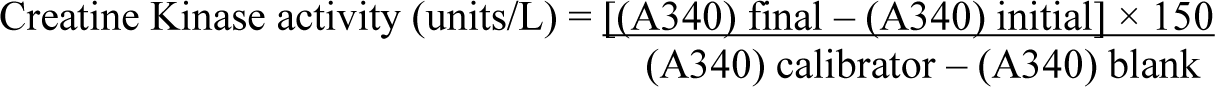

Optimum venom concentration was determined for myotoxicity inhibition assay, based on significant Creatine Kinase activity in serum samples and survival of mice during incubation of 2 hours. An NnV dose of 30 µg was selected for myotoxicity inhibition assay by N194 and N248, based on significant CK level and survival time (Fig. S8A). Similarly, the optimum dose of EcV was estimated to be 40 µg for E10 and E113-mediated myotoxicity inhibition (Fig. S8B).

Ten groups with 5 mice in each group, between weight range of 20-22 grams were used for myotoxicity inhibition assay. Pre-incubated mixtures of venom and different amounts of scFvs were administered in the gastrocnemius muscle of test group mice, as mentioned in table S2B. Vehicle control and positive control groups were injected with saline and venom, respectively. Creatine Kinase activity was estimated for all groups using the same protocol and myotoxicity inhibition was measured for each scFv by comparing it with the venom control group.

### Haemolysis inhibition assay

Human packed RBC were spin washed (1800 rpm; 5 minutes) three times with chilled 1X PBS and resuspended in RPMI (HyClone, GE Healthcare, USA) at 2% haematocrit. A 100 μl suspension was added to a 96-well plate containing the serial dilutions of whole venom at different concentrations (EcV concentration = 2 mg/ml – 0.122 μg/ml; NnV concentration = 5 mg/ml – 0.153 μg/ml). RBC suspension in 100 µl RPMI media alone (for baseline values) and 100 µl of 100% Triton X-100 (SigmaAldrich Corp., USA) with consecutive serial dilutions in RPMI media (for 100% haemolysis) were used as negative control and positive control, respectively. After incubation at 37 °C for 18 hours, the samples were centrifuged, and the supernatant was used to determine the haemolytic activity measured in terms of haemoglobin release as monitored by O.D. at 415 nm (A_415_). The whole venom concentration corresponding to half of the A_415_ value was considered as EC_50_ and the same concentration was further used to assess the effect of scFv mediated haemolysis inhibition.

EC_50_ concentration of whole venom was pre-incubated with several dilutions of different scFvs (EcV concentration = 0.25 mg/ml; NnV concentration = 5 μg/ml) separately and added with washed RBC in different experiments. The same dilutions of scFvs were mixed with RBC and considered as scFv control. Positive and negative control were used as described earlier. All samples were incubated at 37°C for 18 hours and then centrifuged twice at 1800 rpm for 15 minutes. The extent of haemolysis was estimated for all the samples by observing haemoglobin release as monitored by O.D. at 415 nm.

### Screening and crystallization

Attempts were made to crystallize all four venom-neutralizing scFvs and their complexes with respective toxins to get further insights into the structural basis of PLA2 inhibition. Hampton screens were used to screen crystallization conditions. The crystallization droplets of 600 µl size were mixed in 1:1, 1:2, and 2:1 ratio of protein and buffer using Mosquito crystallization robot (TTP-labtech). The plates were kept at 20°C in Rumed incubator. Needle like crystals were obtained in 0.05 M Bis-Tris propane at pH 5.0, 0.05 M Citric acid, and 16% w/v Polyethylene glycol 3,350 by further expansion of crystallization conditions and used for further diffraction. 8 mg/ml protein concentration was used for setting crystallization using hanging-drop vapour diffusion method at 20°C.

### Data collection and structure determination

Crystals were soaked for 5-10 seconds in 15% glycerol as a cryoprotectant and stored in liquid nitrogen for data collection. The crystals of E113 in an apo form showing reasonably good diffraction quality could be obtained, and data were collected at 2.75 Å resolution using the XRD 2 variable energy beamline of Elettra Synchrotron, Trieste (Table S3). iMosflm software was used for data processing. Space group and Laue group were determined using POINTLESS of CCP4 program suite[29]. The log-likelihood gain (LLG) score of 3395 and translation factor Z (TFZ) score of 49.7 signified a good solution towards solving the phase problem. Initial solution of the phase problem was done using PHASER[30] by molecular replacement with the previously published structure of the scFv with PDB ID: 6DSI. Matthews coefficient[31] was calculated from the CCP4 program suite for the number of molecules in the asymmetric unit. Refmac5 of CCP4 software[29] was used for the initial ten rounds of rigid-body refinement and restrained refinement; later Phenix refine program of PHENIX software[32] was used for refinement of the B-factor. Further iterative rounds of refinement were carried out along with model building in COOT[33] (Table S4), with final R_work_ and R_free_ value of 23.26% and 26.41%, respectively. The topology files for ligand refinement was generated using ReadySet program of PHENIX as utilised by Madni *et al.*[34]. PyMOL was used to generate figures. The final quality was checked using the PDB validation server.

### Docking of scFv-toxin complex

To understand the structural basis of scFv mediated PLA2 inhibition, crystal structure of PLA2 (PDB ID: 2QHE) and E113 (PDB ID: 8IA6) were used for molecular docking. Key interacting residues of PLA2 were identified based on known binding regions of PLA2 with its inhibitor suramin in reported PLA2-suramin complex (PDB ID: 3BJW & 1Y4L). Subsequently, the molecular docking was performed of scFv E113 guided by the juxtapositioning of antigen combining site of scFv and the key interacting residues of the PLA2 using the Biologics module of the Schrödinger Suite. The complex was energy minimized using protein preparation wizard[35]. In the protein-protein docking module, various poses were generated. The best docked pose in close proximity to the scFv CDR region and key residues of PLA2 was selected for further analysis. The relative binding affinity of the selected docked pose was also estimated using the Prime module (MM-GBSA) of the Schrödinger Suite[36–38].

### Molecular dynamic simulation

MD simulations were carried out using Desmond module of Schrödinger suite[39]. Docked complex E113/ PLA2 was pre-processed using protein preparation wizard[35]. The system was further optimized by minimization of hydrogen atoms followed by restrained minimization. Furthermore, the system was explicitly solvated using TIP3P with box edges 20 Å from the protein outermost atoms. The system was neutralized by the addition of monovalent counterions Na+ or Cl- and energetically minimized using Desmond module[39]. The production run of 500 ns with OPSL3 force field was subjected for MD simulation[40]. Root mean square deviations (RMSD) and root mean square fluctuations (RMSF) of the trajectories were calculated over the time of 500 ns. Snapshots were extracted at an interval of 100ns and were superimposed in PyMOL to observe the structural variations in the E113/ PLA2 complex.

### Statistical analysis

Statistical significance of the data was estimated using GraphPad PRISM 7 (GraphPad Software, La Jolla California USA, www.graphpad.com) and probability (p) value less than 0.05 were considered significant. Statistical significance of soluble scFv ELISA experiment was evaluated by Sidak’s multiple comparisons test of two-way ANOVA. For survival assay, all scFv treated groups were compared with venom only control groups using Log-rank (Mantel-Cox) test. scFv treated and vehicle control groups were compared with venom challenged group using Dunnett’s multiple comparisons test of one-way ANOVA in myotoxicity inhibition and optimization assay. For haemolysis inhibition and optimization assays, Dunnett’s multiple comparisons test of two-way ANOVA was used to compare venom control with scFv + venom, scFv + RBC and RBC alone.

## Supporting information

PDB validation report

## Acknowledgments

We thank Dr. Deepti Jain and Dr. Vineet Gaur for their useful suggestions. We would like to thank Dr. Asif Mohammed for providing human-packed RBC. We thank Dr. Raghurama Prabhakara Hegde for providing support with data collection at Elettra Sincrotrone Trieste, Italy. We would like to acknowledge the Department of Biotechnology, Govt. of India, for the generous funding.

## Author contributions

D.M.S. and A.K. conceived and designed the experiments. A.K. and Z.K.M. carried out phage display biopanning. A.K. carried out all biochemical assays. A.K. and Z.K.M. determined the crystal structure and carried out computational studies. A.K., Z.K.M., and S.C. carried out animal experiments. A.K., Z.K.M., and D.M.S. analyzed the data. A.K., Z.K.M., and D.M.S. wrote the manuscript. D.M.S. supervised the project. All authors have read and approved the manuscript for publication.

## Conflict of interest

The authors declare no competing interests.

## Ethical approval

Clearance for the use of the animal model was taken from the Institutional Animal Ethics Committee of ICGEB, New Delhi, and IAEC approval no. is ICGEB/IAEC/07032020/SI-17. While using animal models, internationally accepted guidelines and regulations for experimental use and care were followed.

## Funding

We acknowledge the Department of Biotechnology, Government of India, for overall funding and providing fellowship to Amit Kumar.

## Data availability

All data that support the findings of this study are available from the corresponding author upon reasonable request or shared as supplementary information and source files. The atomic coordinates and structure factors have been deposited in the Protein Data Bank, www.wwpdb.org (PDB ID code: 8IA6).

## Abbreviations

CDRs: Complimentary Determining Regions
EcV: *Echis carinatus* venom
NnV: *Naja naja* venom
PhospholipaseA2: PLA2
scFv: single chain Fragment variable

## Notes

https://www.rcsb.org/structure/unreleased/8IA6

### Competing Interest Statement

The authors have declared no competing interest.

