## Supplementary information for "Antibodies against Elapidae and Viperidae snake venoms: in vivo neutralization and mechanistic insights"

**Supplementary Figure 1. Selection of venom-binding scFv phages from human scFv libraries**

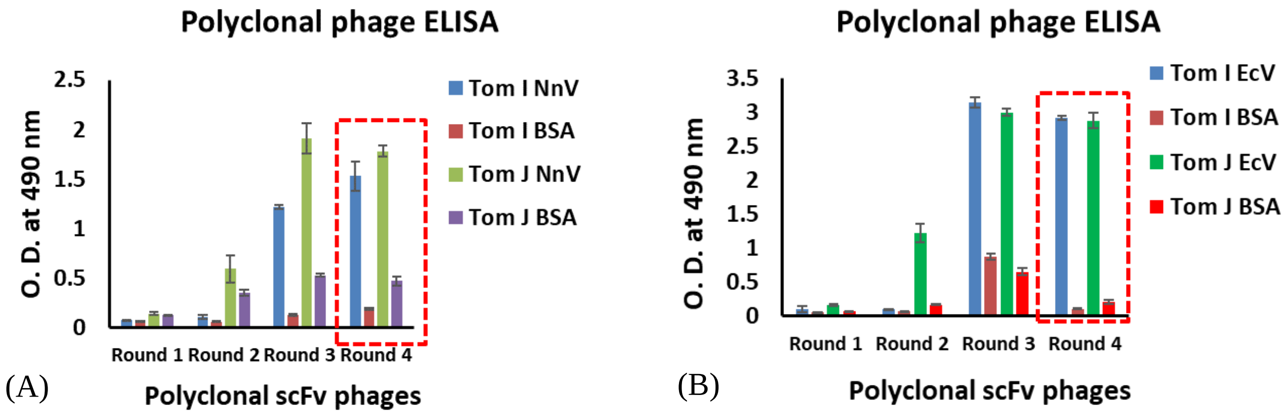


***Fig. S1. Binding analysis of venom-binding polyclonal scFv phages from four rounds of biopanning*** *(A) Binding of Tomlinson I and J polyclonal scFv phages with NnV. (B) Binding of Tomlinson I and J polyclonal scFv phages with EcV. BSA was used as the negative control.*

**Supplementary Figure 2. Purification profile of unique scFvs**

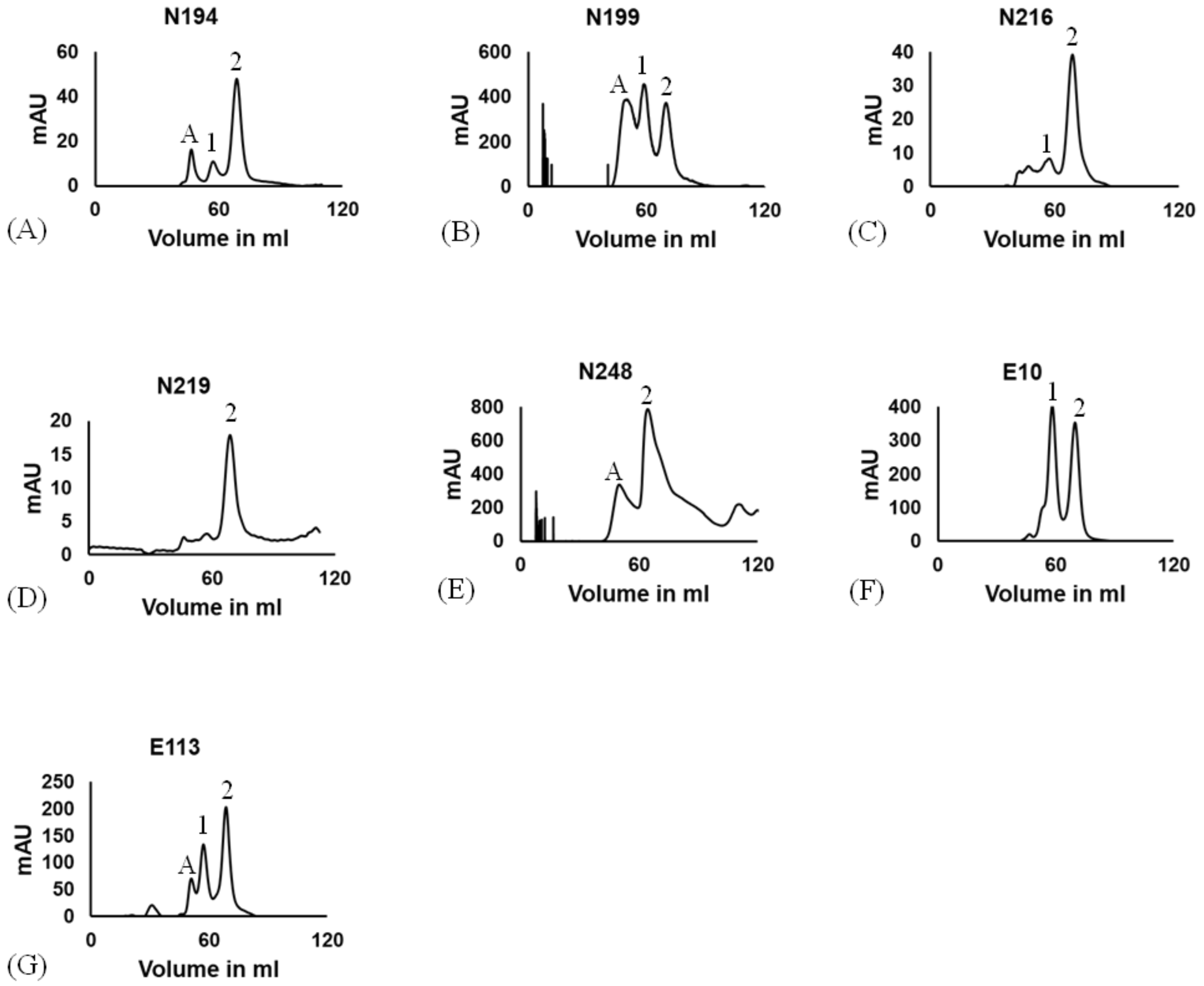


***Fig. S2. Purification of soluble scFv with whole venom.*** *Purification of (A) N194, (B) N199, (C) N216, (D) N219, (E) N248, (F) E10, (G) E113; Peak A, 1, and 2 represent an aggregate, monomeric, and dimeric form of scFvs, respectively.*

**Supplementary Figure 3. Silver stain gels of venom control in the pull-down experiment**

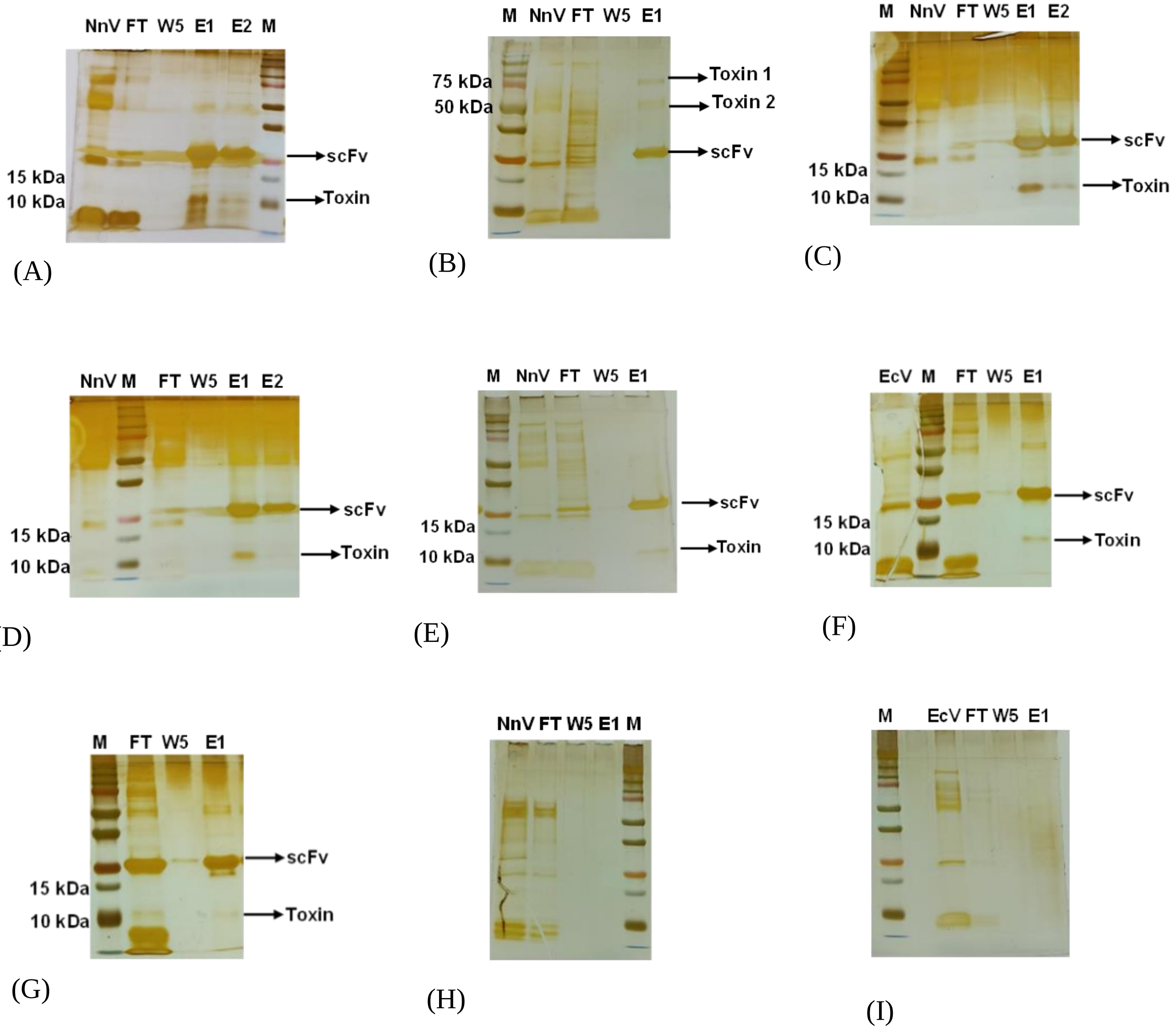


***Fig. S3. Interaction of venom toxins with scFvs.*** *Silver stain gels of (A) N194-toxin complex, (B) N199-toxin complex, (C) N216-toxin complex, (D) N219-toxin complex, (E) N248-toxin complex, (F) E10-toxin complex, (G) E113-toxin complex, (H) NnV negative control, (I) EcV negative control; (M) Protein ladder. Lane M, NnV, EcV, FT, W5, and E1 represent marker, Naja naja whole venom, Echis carinatus whole venom, flow-through, 5th wash fraction, and elute 1, respectively.*

**Supplementary Figure 4. Identification of major toxins in the whole venom**

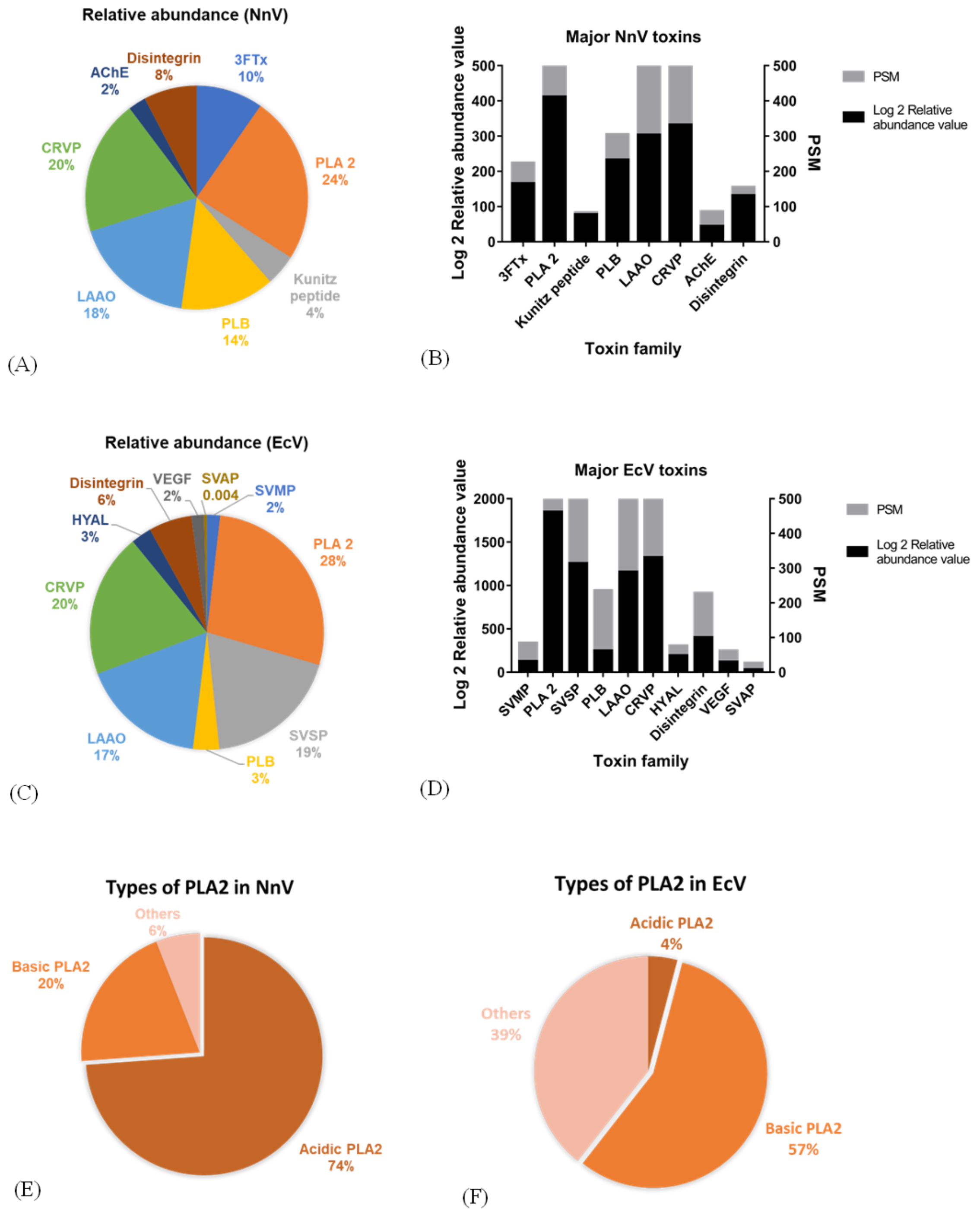


***Fig. S4. Characterization of snake venom proteome.*** *Interspecies and intraspecies variations in the composition of snake venom have been observed, and various factors play a substantial role in diversification*[23–25]*. Therefore, the whole venoms of Naja naja (NnV) and Echis carinatus (EcV) were analyzed for the identification of clinically relevant dominant toxins by mass spectrometry. (A) Percentage relative abundance of the major toxins in NnV (B) Log2 relative abundance value of dominant toxin families of NnV (shown on left Y-axis) with peptide spectrum match (PSM) score match value (shown on right Y- axis) (C) Percentage relative abundance of the major toxins in EcV (D) Log2 relative abundance value of dominant toxin families (shown on left Y-axis) with peptide spectrum match (PSM) value (Shown on right Y-*

*axis) (E) Relative abundance of different PLA2 groups in NnV (F) Relative abundance of different PLA2 groups in EcV (3FTx: Three finger toxin, AChE: Acetylcholinesterase, CRVP: Cysteine-rich venom protein, HYAL: Hyaluronidase, LAAO: L-amino-acid oxidase, PLA2: Phospholipase A2, PLB: Phospholipase B, SVAP: Snake venom aspartic proteases, SVMP: Snake venom metalloproteases, VEGF: Vascular endothelial growth factor)*

### Table S1. Venom and scFvs doses for MLD estimation and survival assay

| | | |
|---|---|---|
| **A. Minimum lethal dose estimation** | | |
| Group number | NnV dose (in mg/kg of mice body weight) | EcV dose (in mg/kg of mice body weight) |
| Group 1 | Phosphate buffer saline (Control) | Phosphate buffer saline (Control) |
| Group 2 | 0.2 | 0.5 |
| Group 3 | 0.25 | 0.55 |
| Group 4 | 0.3 | 0.6 |
| Group 5 | 0.35 | 0.75 |
| Group 6 | 0.4 | 1 |
| Group 7 | 0.45 | 1.25 |
| Group 8 | 0.5 | 1.5 |
| Group 9 | 0.75 | Not used |
| Group 10 | 1 | Not used |
| **B. scFv dose optimization for *in vivo* survival assay** | | |
| | EcV challenged mice | |
| Group 1 | Phosphate buffer saline (Vehicle control) | |
| Group 2 | 3(MLD) EcV (Venom only control) | |
| Group 3 | 2 mg E10 + 3(MLD) EcV | |
| Group 4 | 4 mg E10 + 3(MLD) EcV | |
| Group 5 | 2 mg E113 + 3(MLD) EcV | |
| Group 6 | 4 mg E113 + 3(MLD) EcV | |
| Group 7 | 1 mg E10 + 1 mg E113 + 3(MLD) EcV | |
| Group 8 | 2 mg E10 + 2 mg E113 + 3(MLD) EcV | |
| Group 9 | Anti EcV serum + 3(MLD) EcV | |
| **C. *In vivo* survival assay** | | |
| | NnV challenged mice | EcV challenged mice |
| Group 1 | Phosphate buffer saline (Vehicle control) | Phosphate buffer saline (Vehicle control) |
| Group 2 | 2× MLD EcV (Venom only control) | 2(MLD) EcV (Venom only control) |
| Group 3 | 0.5 mg N194 + 2(MLD) EcV | 0.5 mg E10 + 2(MLD) EcV |
| Group 4 | 1 mg N194 + 2(MLD) EcV | 1 mg E10 + 2(MLD) EcV |
| Group 5 | 2 mg N194 + 2(MLD) EcV | 2 mg E10 + 2(MLD) EcV |
| Group 6 | 0.5 mg N248 + 2(MLD) EcV | 0.5 mg E113 + 2(MLD) EcV |

| | | |
|---|---|---|
| Group 7 | 1 mg N248 + 2(MLD) EcV | 1 mg E113 + 2(MLD) EcV |
| Group 8 | 2 mg N248 + 2(MLD) EcV | 2 mg E113 + 2(MLD) EcV |
| Group 9 | 0.25 mg N194 + 0.25 mg N248 + 2(MLD) EcV | 0.25 mg E10 + 0.25 mg E113 + 2(MLD) EcV |
| Group 10 | 0.5 mg N194 + 0.5 mg N248 + 2(MLD) EcV | 0.5 mg E10 + 0.5 mg E113 + 2(MLD) EcV |
| Group 11 | 1 mg N194 + 1 mg N248 + 2(MLD) EcV | 1 mg E10 + 1 mg E113 + 2(MLD) EcV |
| Group 12 | Anti-NnV serum + 2(MLD) EcV | Anti-EcV serum + 2(MLD) EcV |

**Supplementary Figure 5. Minimum lethal dose of NnV and EcV**

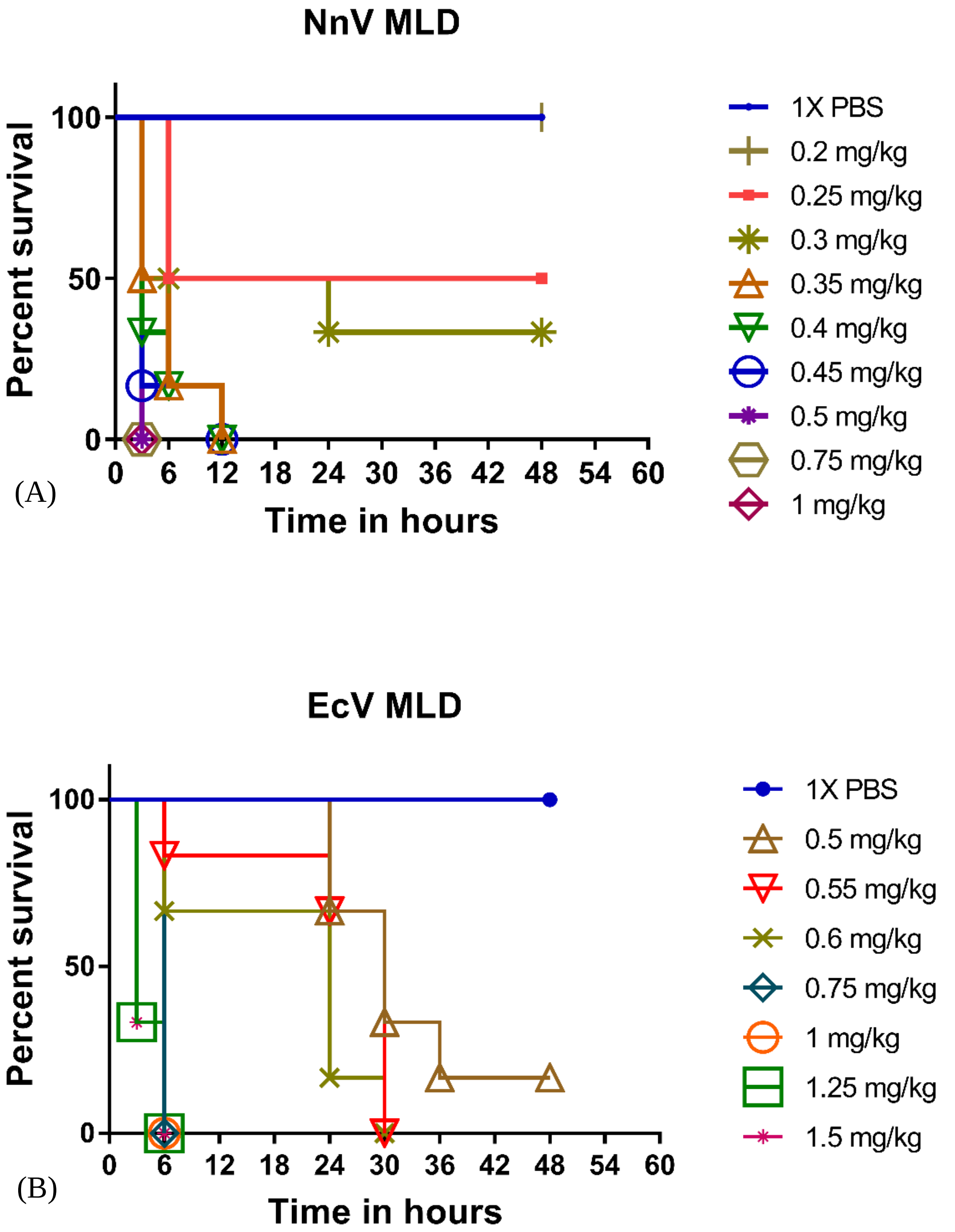


***Fig. S5. In vivo minimum lethal dose estimation*** *(A) NnV, and (B) EcV; for all groups n=6 and p<0.05 when vehicle control groups were compared with mice groups challenged with NnV dose of 0.3 to 1 mg/kg of mice body weight or any EcV dose.*

**Supplementary Figure 6. Binding of serum sample with *Naja naja* and *Echis carinatus* venom after immunization of BALB/c mice**

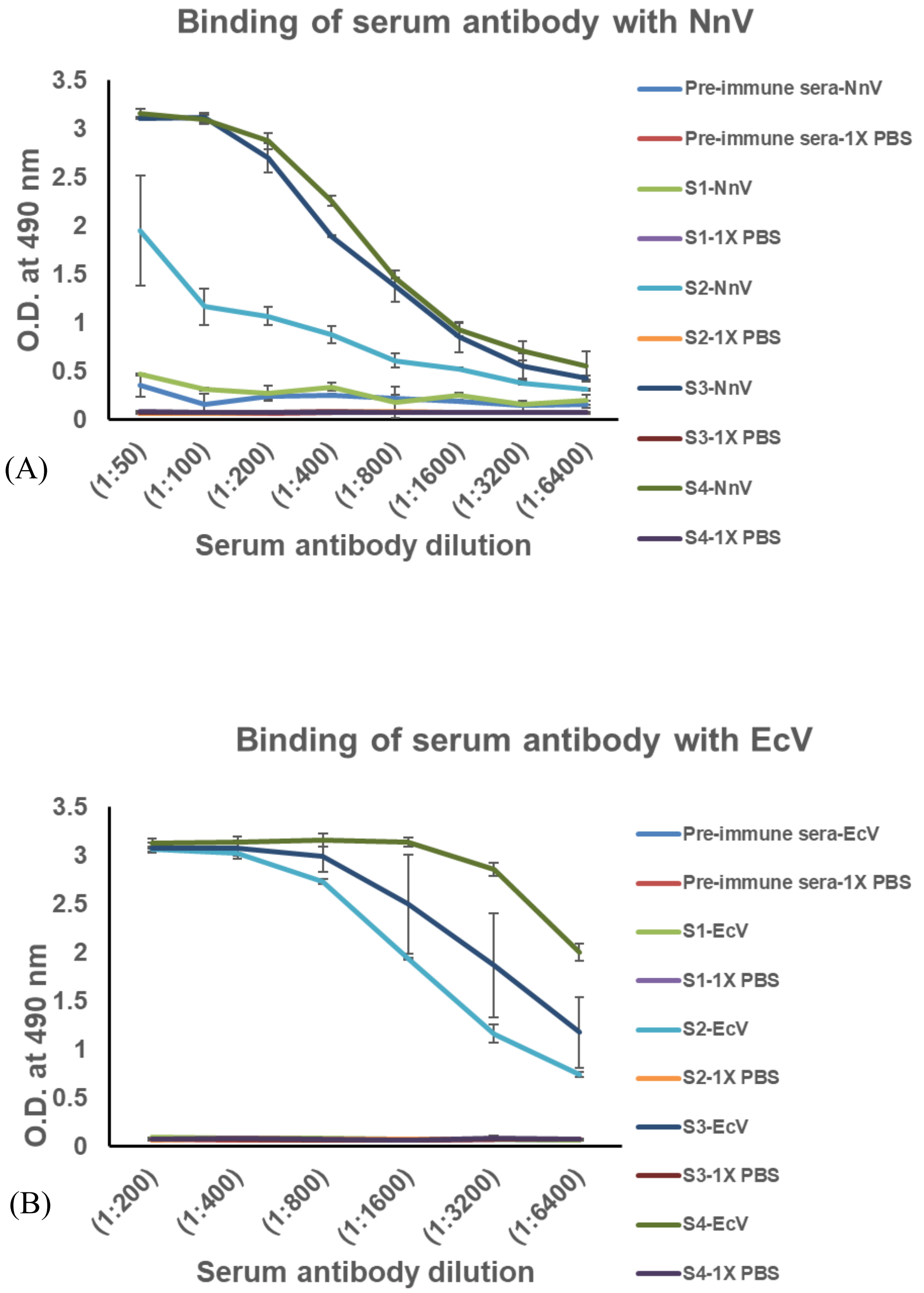


***Fig. S6. Binding of serum samples with whole venom*** *(A) Anti-NnV serum with NnV, and (B) Anti-EcV serum with EcV; for all groups n=3 (S1- sera sample collected after 10 days of first immunization, S2- sera sample collected after 10 days of first booster immunization, S3- sera sample collected after 10 days of second booster immunization, S4- S2- sera sample collected after 10 days of third booster immunization)*

**Supplementary Figure 7. Optimization of scFv dose for *in vivo* protection assay**

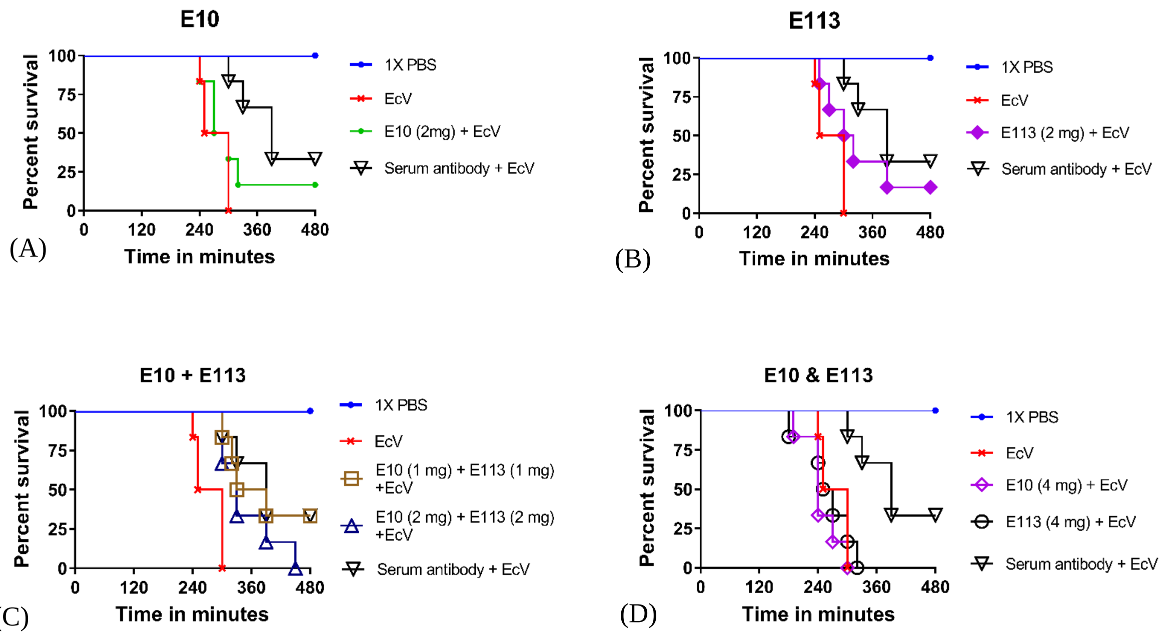


***Fig. S7. Anti-PLA2 scFvs mediated in vivo protection from the whole venom of Echis carinatus.*** *Different groups of mice were intraperitoneally administered with 3xMLD of EcV. Survival effects of respective scFvs were evaluated in the mice groups which were given intraperitoneal injections of pre-incubated mixtures of 3xMLD of EcV and different doses of scFvs. Kaplan-Meier survival analysis in (A) E10 treated group. (B) E113 treated group. The E10 and E113 demonstrated 16% survival in mice challenged with 3x(MLD) of EcV in an 8-hour survival experiment. (C) The groups were treated with co-administration of 1 mg E10 and 1 mg E113. Combined administration of a 2 mg dose, comprising an equal quantity of both scFvs, resulted in enhanced survival of 33%. (D) The groups were treated with co-administration of 2 mg E10 and 2 mg E113. 4 mg dose comprising individual scFv or a mixture of both scFvs, resulted in a decline in survival percentage. (Vehicle control, venom-only control, and serum antibody control groups were same for all four graphs A, B, C, and D. Survival experiments pertaining to graphs A, B, C, and D. were performed simultaneously.) Each individual curve indicates the survival status of a group of 6 mice. $p<0.05$ when venom-challenged groups were compared with the mice groups administered with pre-incubated mixtures of 3xMLD of EcV and (1 mg E10 + 1 mg E113) or (2 mg E10 + 2 mg E113) or serum antibody.*

### Table S2. Venom and scFv dose for myotoxicity inhibition assay

| **A. Optimum dose estimation for myotoxicity** | | |
|---|---|---|
| Group number | NnV dose | EcV dose |
| Group 1 | Phosphate buffer saline (Control) | Phosphate buffer saline (Control) |
| Group 2 | 20 µg | 20 µg |
| Group 3 | 30 µg | 30 µg |
| Group 4 | 40 µg | 40 µg |
| Group 5 | 50 µg | 50 µg |
| **B. Myotoxicity inhibition assay** | | |
| | NnV challenged mice | EcV challenged mice |
| Group 1 | Phosphate buffer saline (Vehicle control) | Phosphate buffer saline (Vehicle control) |
| Group 2 | 30 µg NnV (Venom only control) | 40 µg EcV (Venom only control) |
| Group 3 | 0.2 mg N194 + 30 µg NnV | 0.2 mg E10 + 40 µg EcV |
| Group 4 | 0.4 mg N194 + 30 µg NnV | 0.4 mg E10 + 40 µg EcV |
| Group 5 | 0.6 mg N194 + 30 µg NnV | 0.6 mg E10 + 40 µg EcV |
| Group 6 | 0.8 mg N194 + 30 µg NnV | 0.8 mg E10 + 40 µg EcV |
| Group 7 | 0.2 mg N248 + 30 µg NnV | 0.2 mg E113 + 40 µg EcV |
| Group 8 | 0.4 mg N248 + 30 µg NnV | 0.4 mg E113 + 40 µg EcV |
| Group 9 | 0.6 mg N248 + 30 µg NnV | 0.6 mg E113 + 40 µg EcV |
| Group 10 | 0.8 mg N248 + 30 µg NnV | 0.8 mg E113 + 40 µg EcV |

**Supplementary Figure 8. Optimization of venom control for myotoxicity inhibition assay**

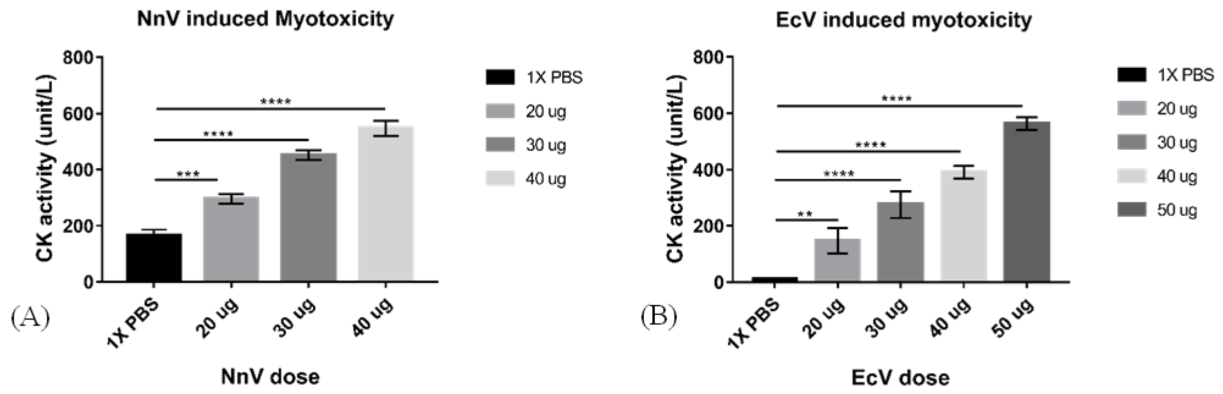


***Fig. S8. Myotoxicity estimation in mice challenged by whole venom of Echis carinatus and Naja naja:*** *Myotoxicity was estimated by measuring Creatine Kinase activity in the serum samples of mice injected (intramuscular injection in gastrocnemius muscle) with different doses of whole venom. Creatine Kinase activity in serum sample of (A) NnV-challenged mice (B) EcV-challenged mice. Each data point represents the mean of triplicates, and error bars indicate standard deviations from the mean of each data set. For all the graphs n=3 and p<0.05 (**p= 0.0021, ***p=0.0002, ****p<0.0001).*

**Supplementary Figure 9. Optimization of optimum dose of whole venom control for hemolysis inhibition assay**

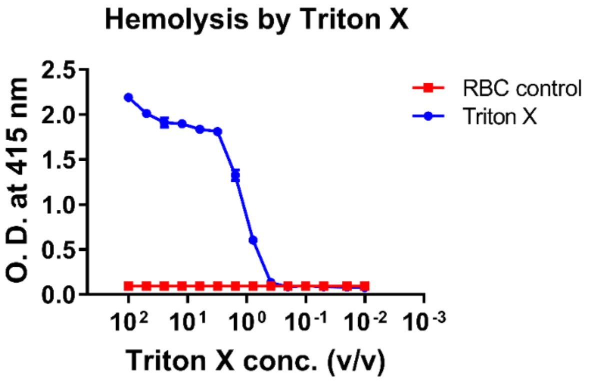


(A)

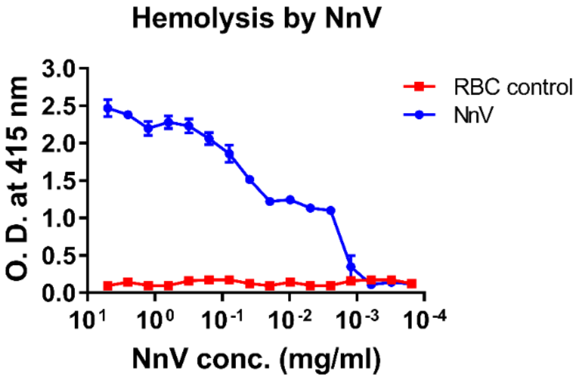


(B)

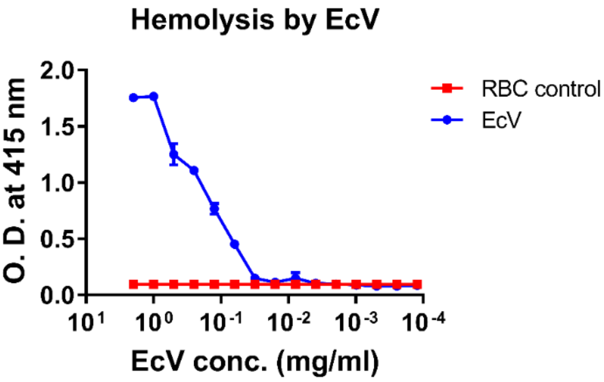


(C)

***Fig. S9. Estimation of human packed RBC lysis by*** *(A) Triton X, (B) NnV, (C) EcV; $p<0.05$ when RBC control is compared with RBC lysed by Triton X, NnV or EcV*

**Table S3. : Data collection statistics**

| | **8IA6** |
|---|---|
| X-ray source | Elettra Beamline (XRD 2) |
| Detector | Pilatus 6M |
| Space group | *I* 121 |
| Cell dimensions | |
| *a, b, c* (Å) | 66.88, 53.87, 169.32 |
| α, β, γ | 90.00, 94.42, 90.00 |
| Resolution (Å) | 2.75 |
| $R_{p.i.m}$ | 16.4 (35.8) |
| *CC1/2* (%) | 92.5 (61.3) |
| $I / \sigma I$ | 4.2 (2.1) |
| Completeness (%) | 95.37 (98.23) |
| Redundancy | 2.8 (2.7) |

--------------------------------------------

*The values in parentheses refer to the highest-resolution shell.

**Table S4. : Refinement statistics (molecular replacement)**

| | **8IA6** |
|---|---|
| Resolution (Å) | 29.23 – 2.75 |
| No. reflections | 15192 (1556) |
| $R_{work}$ / $R_{free}$ | 0.2326 / 0.2641 |
| No. of atoms | |
| Protein | 3372 |
| Ligand/ion | 57 |
| Water | 165 |
| Average B-factor | 33.36 |
| R.m.s. deviations | |
| Bond lengths(Å) | 0.003 |
| Bond angles (°) | 0.67 |
| Ramachandran statistics | |
| Favoured (%) | 95.41 |
| Allowed (%) | 4.13 |
| Outliers (%) | 0.46 |

----------------------------------------

*The values in parentheses refer to the highest-resolution shell.

**Supplementary Figure 10. Stability of scFv-113-α-EcV/S49-PLA2 complex**

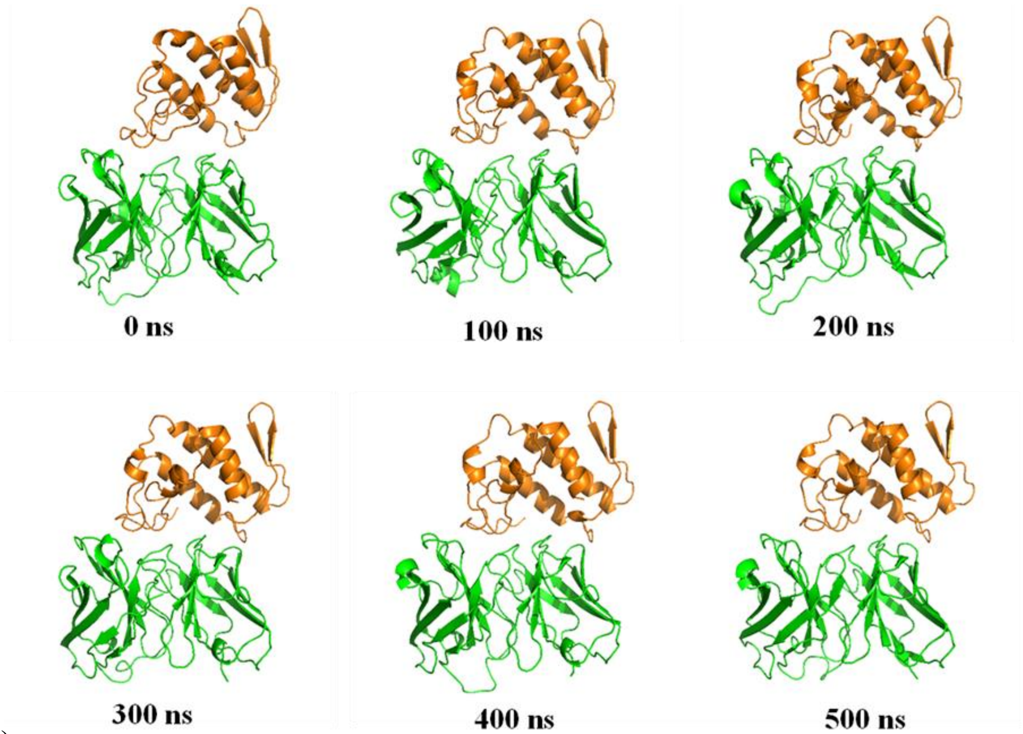


(A)

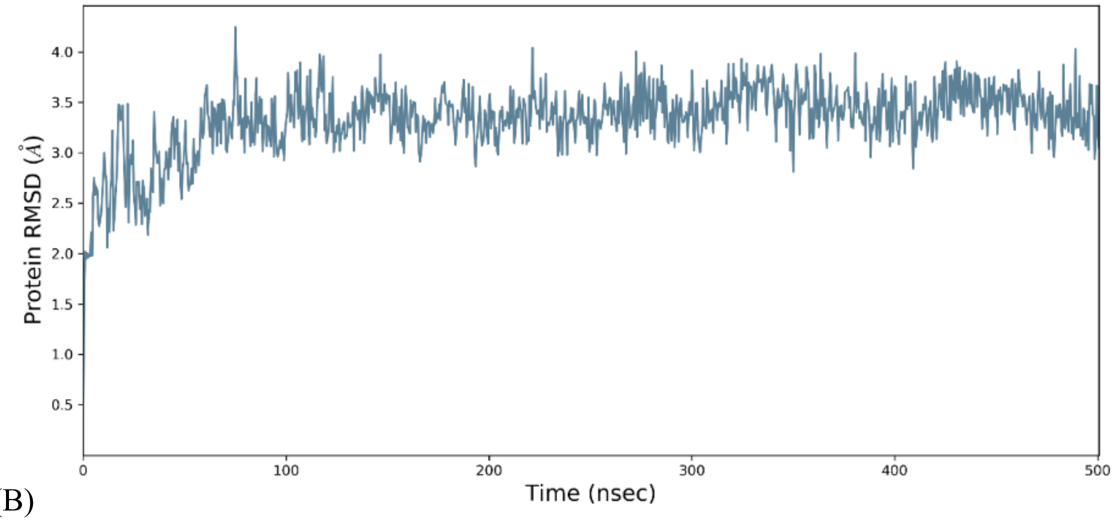


(B)

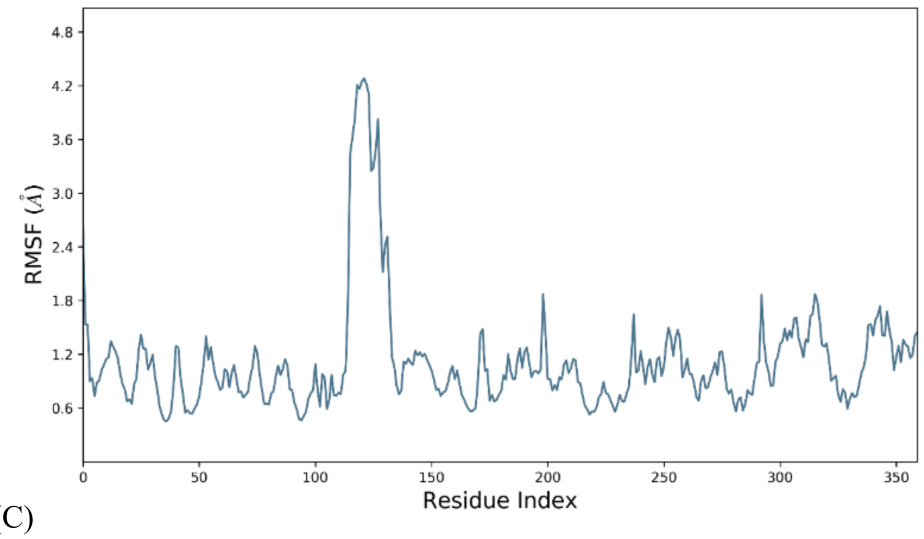


(C)

***Fig. S10. Comparison of modelled structures of scFv-113-α-EcV/S49-PLA2 complex during the simulation run.*** *(A) The PDBs of the complex were extracted at 100 ns intervals during the 500 ns run. E113 was represented in green, and S49-PLA2 was represented in orange. (B) Root Mean Square Deviation of E113/S49-PLA2 complex for 500 ns. (C) Root Mean Square Fluctuation of each residue of E113/S49-PLA2*
